# Rice MEL2 regulates the timing of meiotic transition as a component of cytoplasmic RNA granules

**DOI:** 10.1101/2021.03.24.433842

**Authors:** Manaki Mimura, Seijiro Ono, Ken-Ichi Nonomura

## Abstract

Cytoplasmic RNA granules play important roles in gene expression at the post-transcriptional level. In this study, we found that the rice RNA-binding protein MEIOSIS ARRESTED AT LEPTOTENE2 (MEL2), which contributes to the control of meiotic entry timing, was a constituent of RNA granules, frequently associating with processing bodies and stress granules in the cytoplasm of premeiotic spore mother cells. MEL2 has four conserved domains and a large intrinsically disordered region, which is often responsible for formation and maintenance of granular structures. MEL2-like proteins with diverse domain structures are widely conserved in land plants and charophyte algae. In basal land plants, MEL2-like proteins are exclusively expressed in the sporophyte, which expresses meiotic genes, suggesting the functional conservation of MEL2 among land plant species. We propose here that MEL2 participates in post-transcriptional regulation of meiotic genes as a component of RNA granules to ensure proper timing of the meiotic transition.

## Introduction

Meiosis is a special type of cell division that produces haploid gametes through two rounds of chromosome segregation without intervening DNA replication. Prior to meiosis, cells that have decided to enter the meiotic process are irreversibly committed to the completion of meiosis [1]. After commitment, cells undergo meiosis-specific DNA replication, called premeiotic DNA replication or premeiotic S phase.

Strict control of the mitosis-to-meiosis transition played essential roles in land plant evolution. Charophyte algae, from which ancient land plants had originated [2, 3], undergo zygotic meiosis [4], in which the zygote initially divides by meiosis after karyogamy. By contrast, in all land plant species, zygotic cells proliferate by intercalated mitosis before meiosis [5]. These intercalated mitoses produce a multicellular sporophyte, resulting in an increase in cell numbers prior to entry into the meiotic pathway [6]. In other words, in all land plants, the transition to meiosis is repressed and delayed until sporophytic development is completed. Although many factors involved in maintaining the sporophyte have been identified [7], the mechanisms that directly repress and delay meiosis during the sporophytic phase have largely remained elusive.

In angiosperms, the first germ cell founders, called archesporial cells, are initiated at the hypodermis of the stamen and ovule primordia, which are the male and female organs, respectively. Archesporial cells divide mitotically and differentiate into sporogenous cells, and the surrounding somatic cells are specified for the nursery of interior sporogenous cell lineages. Many factors required for the meiotic cell fate decision and initiation of meiosis have been identified in model plants. The rice Argonaute protein MEIOSIS ARRESTED AT LEPTOTENE1 (MEL1), which is required for normal meiosis, begins to be expressed in archesporial initial cells [8, 9], indicating that these cells are already destined to acquire a meiotic fate. OsSPL, orthologous to the *Arabidopsis* transcription factor SPOROCYTELESS/NOZZLE (SPL/NZZ) [10, 11], plays essential roles in establishment of sporogenous cell identity and meiotic fate acquisition [12]. In *Arabidopsis* female meiocytes, specific downregulation of stem cell factor WUSCHEL (WUS) by RETINOBLASTOMA-RELATED1 (RBR1) is necessary for meiotic cell specification and normal entry to meiosis [13]. *Arabidopsis* SWITCH1 (SWI1), the ortholog of maize AM1, which was initially identified as a key factor in meiotic entry, protects sister chromatid cohesion from the cohesin dissociation factors WAPL from premeiotic S phase to early meiotic prophase I [14–16]. In addition, redox status in anther lobes is important for cell fate specification as well as meiotic entry [17–19]. Thus, the regulation of meiotic initiation has been extensively studied from the standpoints of meiotic cell fate acquisition and meiotic chromosome structures. Nevertheless, it remains obscure how plant sporogenous cells leave the mitotic cell cycle to enter meiosis.

Another unresolved issue in the sexual reproduction of flowering plants is the mechanism by which male meiosis occurs synchronously (on average) among a large number of pollen mother cells (PMCs) in the same anther locule [20]. We previously identified the rice gene *MEL2*, in which loss of function disrupted the synchrony of male meiosis [21]. In *mel2* mutant anthers, most PMCs aberrantly continue mitosis, and a small subset enters meiosis but with delayed timing. These observations indicate that MEL2 is involved in modulating the proper timing of meiotic entry.

MEL2 is an RNA-binding protein that accumulates in the cytoplasm of premeiotic PMCs in rice anthers [21, 22], prompting us to speculate that it is a component of the cytoplasmic ribonucleoprotein (RNP) granule implicated in post-transcriptional regulation. Post-transcriptional regulation is important for meiotic entry in many organisms. For example, in fission yeast, nitrogen starvation promotes the expression of the RNA-binding protein Mei2, which forms dot structures with non-coding RNA on the *Sme2* locus to trap and inactivate the machinery for meiotic mRNA elimination, causing a switch from the mitotic expression pattern to the meiotic pattern [23]. The mouse DExH-box RNA helicase YTHDC2, which is associated with RNA granules, binds *Cyclin A2* mRNA to regulate proper switching from mitosis to meiosis [24]. Deleted in Azoospermia (DAZ) and DAZ-like (DAZL) proteins also play key roles in germ cell development [25]. *Drosophila* Boule, a DAZL family member, binds the 3’-UTR of *Twine* mRNA, which encodes a meiotic type of CDC25 phosphatase, and promotes the premeiotic G2/M transition [26].

In this study, we found that rice MEL2 is a component of cytoplasmic RNA granules in PMCs at premeiotic stages. Our analysis revealed that MEL2-like proteins with diverse domain combinations are widely conserved among land plant species. Our findings demonstrate that the cytoplasmic RNP granule plays an essential role in posttranscriptional regulation of proper timing of meiotic entry in plants.

## Results

### MEL2 is required for proper meiotic gene expression

To assess the impact of MEL2 function on meiotic gene regulation, we performed mRNA-seq analysis of wild-type and *mel2* mutant anthers at three different meiotic stages: premeiosis_early (PE), premeiosis_late (PL), and meiosis_early (ME) (see Supplementary Table 1 for stage definitions).

The level of *MEL2* transcription was highest in PE, followed by PL and ME, when the expression of eleven well-studied meiotic genes peaked (Fig. 1). Hundreds of protein-coding transcripts were differentially expressed in *mel2* mutant anthers, and many of them, including the aforementioned meiotic genes, were down-regulated (Fig. 1, Supplementary Fig. 1a). Gene Ontology (GO) analysis of genes down-regulated in the *mel2* mutant revealed that GO terms associated with meiosis were highly enriched in PL and ME (Supplementary Fig. 1b). These results indicate that the MEL2 function is essential for shifting the transcriptomic landscape to the meiotic pattern in premeiotic rice anthers.

**Fig. 1.**
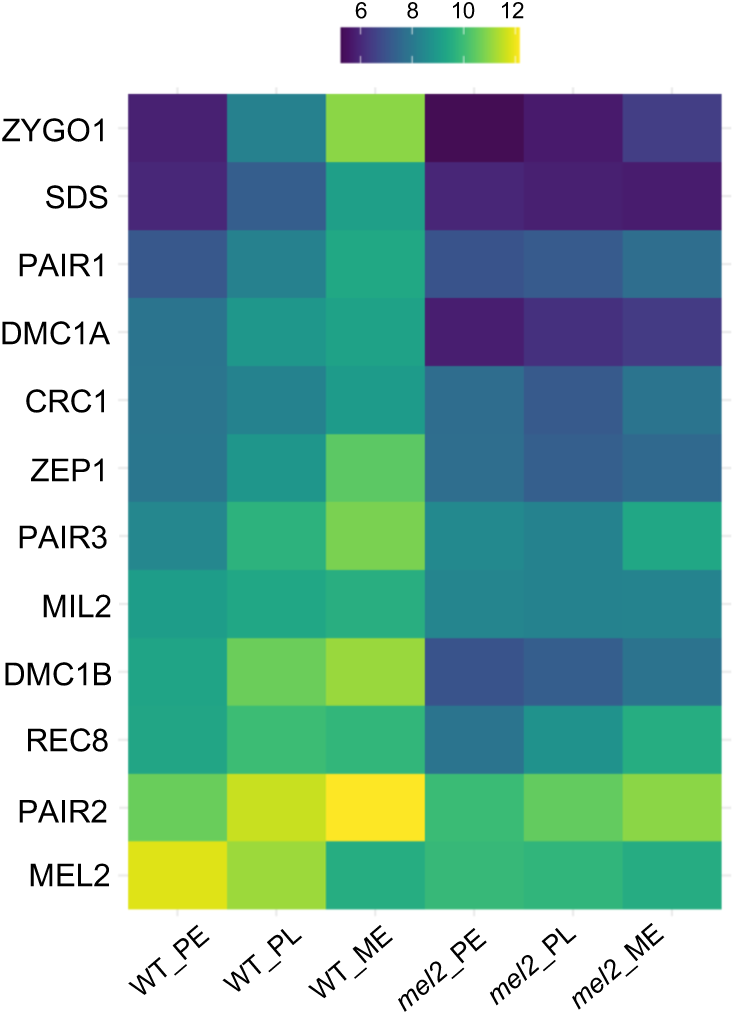
MEL2 is required for normal meiotic gene expression. Heatmap of *MEL2* and 11 meiotic gene transcripts in wild-type and *mel2* mutant anthers at premeiosis and early meiosis. MEL2 expression peaked prior to meiotic gene expression. PE: Pre-meiotic early stage, PL: Pre-meiotic late stage, ME: Meiosis early stage.

### MEL2 takes part in granule formation in the cytoplasm of premeiotic PMCs

Next, we introduced the MEL2-GFP transcriptional fusion construct driven by the MEL2 promoter into *mel2* homozygous plants. The transgene rescued the sterile *mel2* phenotype, indicating that MEL2-GFP was functional *in vivo* (Supplementary Fig. 2).

In PE and PL anthers, MEL2-GFP formed distinct granular foci in addition to diffuse signals in the PMC cytoplasm (Fig. 2a, b). The signals were prominent at PMCs, but not at surrounding anther somatic cells. By contrast, few MEL2-GFP signals were detected in PMCs of ME anthers, which are characterized by meiotic chromosomes with condensed filamentous structures (Fig. 2c). MEL2-GFP expression was also detected in the cytoplasm of megaspore mother cells in developing ovules, but not in surrounding somatic cells (Supplementary Fig. 3), indicating that MEL2 is functional at premeiotic stages in both male and female germ cells.

**Fig. 2.**
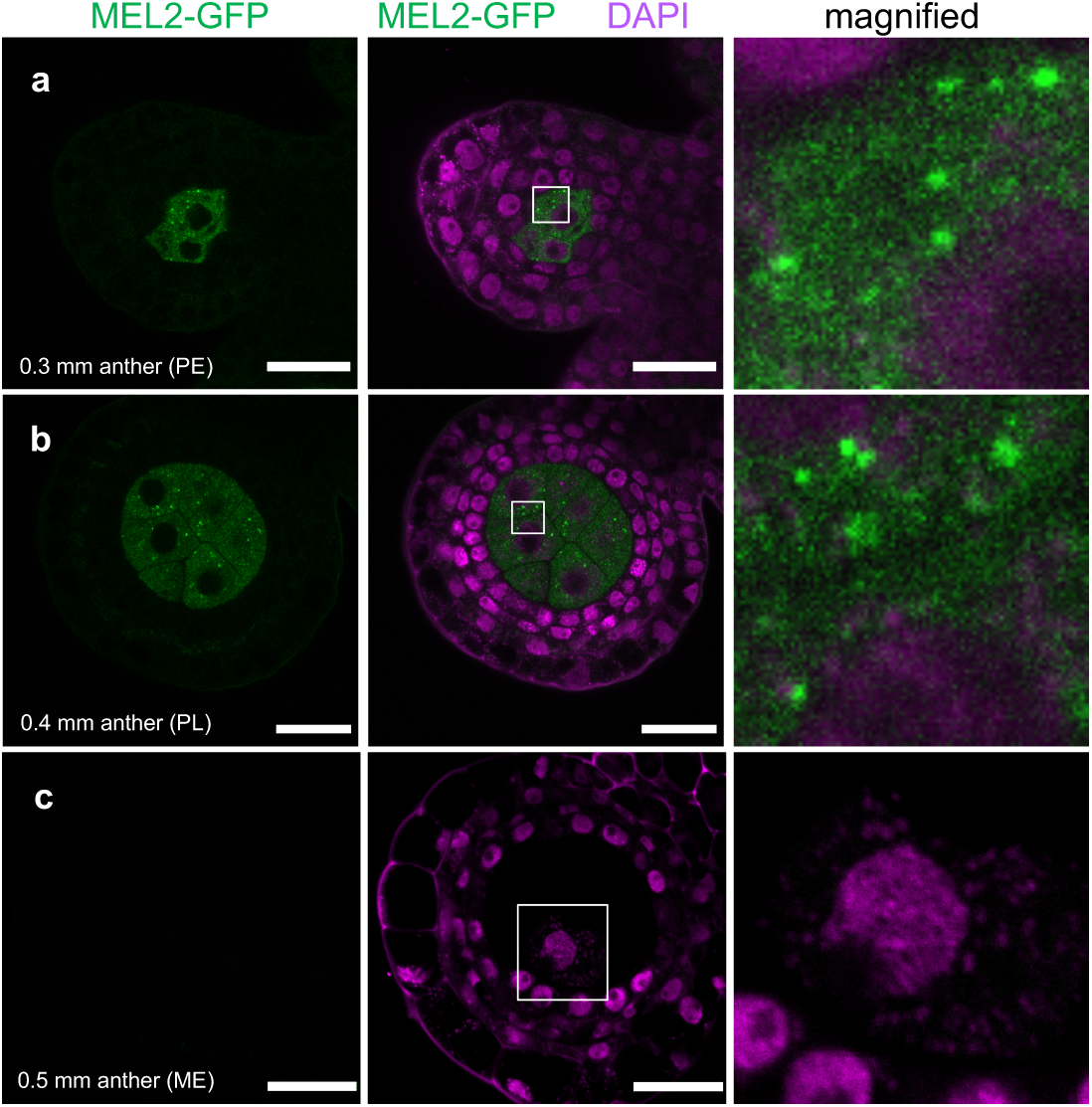
Cytoplasmic MEL2 granules in PMCs of rice premeiotic anthers. Representative confocal images of MEL2-GFP localization in cross-sections of (**a**) PE-, (**b**) PL-, and (**c**) ME-stage anther lobes in transgenic rice plants. The rightmost panels are magnified views of white squares in the middle panels. Bars = 20 μm.

Next, to scale the timing of MEL2 granule formation with the occurrence of premeiotic S, we observed MEL2-GFP signals along with immunofluorescence of PAIR2, a rice HORMA protein required for homolog synapsis [27]. Prior to meiosis, PAIR2 starts to accumulate at the nucleoplasm of germ cells, just as it does following premeiotic S initiation. In PE-stage anthers containing PMCs with faint and diffuse PAIR2 signals in nuclei, indicating that the cells were in the process of undergoing the premeiotic G1/S transition, we observed a few MEL2-GFP granules in the PMC cytoplasm (Fig. 3a). In PL-stage PMCs with more intense diffuse PAIR2 signals in nuclei, indicating cells around premeiotic S/G2, MEL2 granules became slightly but significantly more numerous and larger in volume (Fig. 3b, d–f). The gradual increase in numbers and volumes during premeiotic stage progression indicates that the observed granule formation represents natural MEL2 behavior and was not due to artifacts of anther fixation. After the cells entered meiotic prophase I, characterized by filamentous PAIR2 signals on chromosomes, no MEL2 signal was detected (Fig. 3c). These results confirmed that MEL2 has the potential to form granular structures in the PMC cytoplasm prior to or during premeiotic S phase, but is dispensable for early meiotic events.

**Fig. 3.**
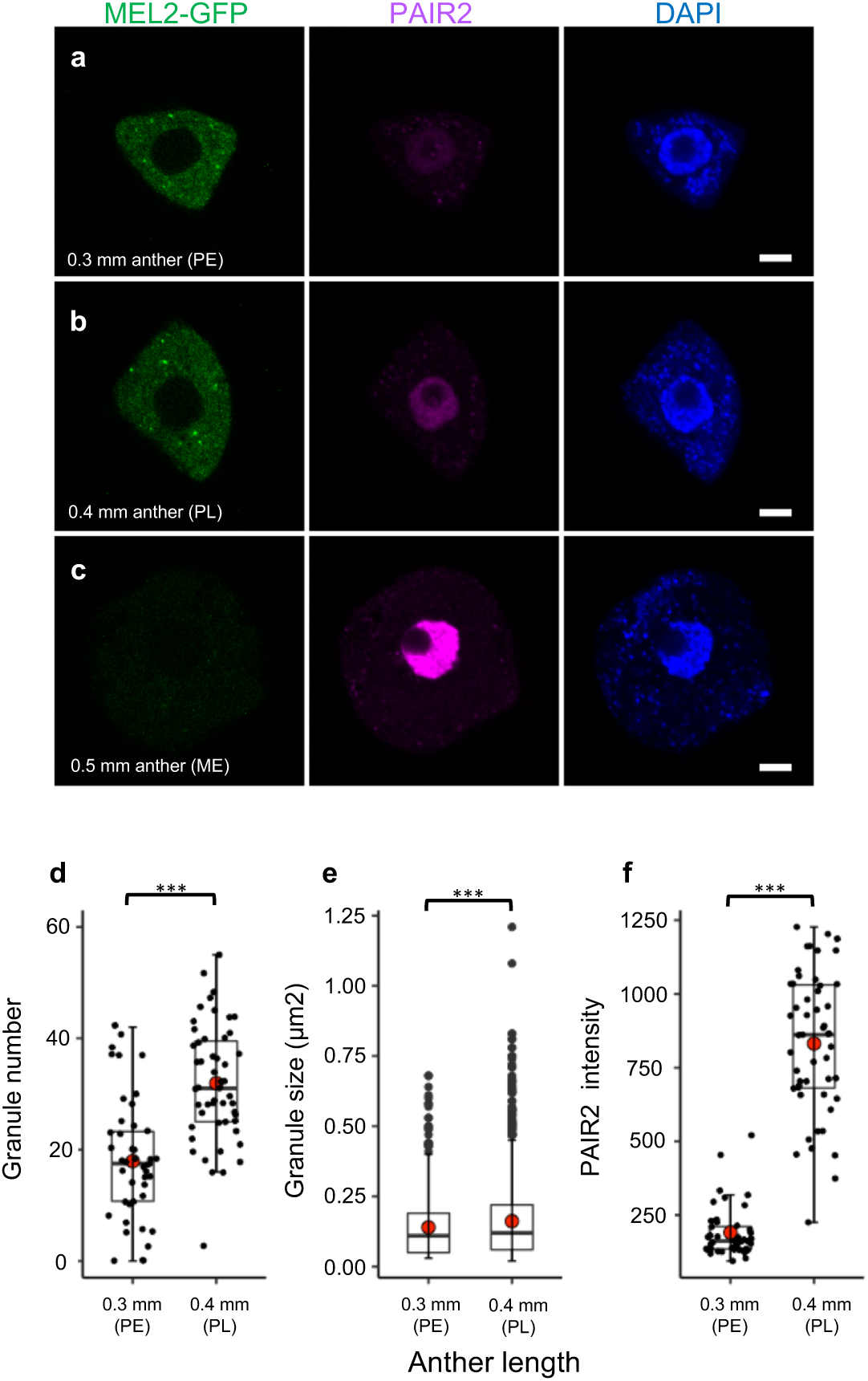
Behavior of MEL2 granules in PMCs around premeiotic S phase. (**a–c**) Simultaneous detection of MEL2-GFP and PAIR2 protein in PMCs at the PE (**a**), PL (**b**), and ME stages (**c**). Granular and diffuse MEL2 signals were detected in the cytoplasm of PMCs at the onset of PAIR2 accumulation in the nucleoplasm (**a, b**), but the signals disappeared by meiotic prophase I (zygotene), when PAIR2 was loaded onto chromosomes (**c**). Bars = 5 μm. (**d–f**) Box plots of the number (**d**) and size (**e**) of MEL2 granules, and mean values of PAIR2 intensity (**f**) in 44 PMCs in PE anthers (0.3 mm) and 51 PMCs in PL anthers (0.4 mm). Red circles indicate averages of plotted values (closed circles). For granule size, the areas of 881 and 1648 granular foci in the aforementioned 44 and 51 PMCs, respectively, were measured and plotted. Triple asterisks indicate a statistically significant difference compared between 0.3 mm (PE) and 0.4 mm (PL) anthers (*t*-test, P < 0.001).

### MEL2 has the ability to form granules with stress-granule components

The results described above raised the possibility that premeiotic and cytoplasmic MEL2-GFP aggregates are equivalent to RNP granules. To test this idea, we co-expressed MEL2-GFP with mCherry (mCh)-tagged components of known RNP granules in rice protoplast cells; the transcription of all constructs was driven by the constitutively active promoter of the rice *Actin* gene. In most cells, MEL2-GFP granular structures were prominent in the cytoplasm, with no diffuse signals (Supplementary Fig. 4a), likely due to aggregation of excess MEL2 due to over-transcription. When co-expressed with mCh-tagged OsDCP1 (Os12g0156400), a rice P-body component involved in mRNA decapping, MEL2 granules frequently adhered with OsDCP1 signals in the cytoplasm (Supplementary Fig. 4b). In the case of OsUBP1b (Os11g0620100), a stress-granule component, a few MEL2 granules were colocalized with UBP1 foci in the cytoplasm, although in cells cultured at 28°C, UBP1-mCh signals were prominent at the nucleus and diffuse in cytoplasm (Supplementary Fig. 4c). Interestingly, when the cells were cultured at 45°C, the colocalized OsUBP1b and MEL2-GFP aggregates in the cytoplasm were highly enlarged (Supplementary Fig. 4d).

To determine whether these observations would be reproducible in premeiotic anthers, we produced transgenic rice plants co-expressing MEL2-GFP with OsDCP1-mCh or OsUBP1b-mCh; transcription of all constructs was driven by the corresponding genes’ promoters (Supplementary Fig. 5a). In anthers, including PL-stage meiocytes, OsDCP1-mCh signals were prominent in the cytoplasm of somatic layered cells (Fig. 4a, Supplementary Fig. 5b), and faint signals were detected in PMCs. In PMCs, 30.0% and 37.2% of 43 OsDCP1-mCh foci completely and partially overlapped with MEL2-GFP foci, respectively (Fig. 4a, d, Supplementary Fig. 5b). In contrast to MEL2, P-bodies were preserved in the PMC cytoplasm at the ME stage (Supplementary Fig. 6). These results suggest that cytoplasmic MEL2 granules are distinct from, but interact with, P-bodies in premeiotic PMCs.

**Fig. 4.**
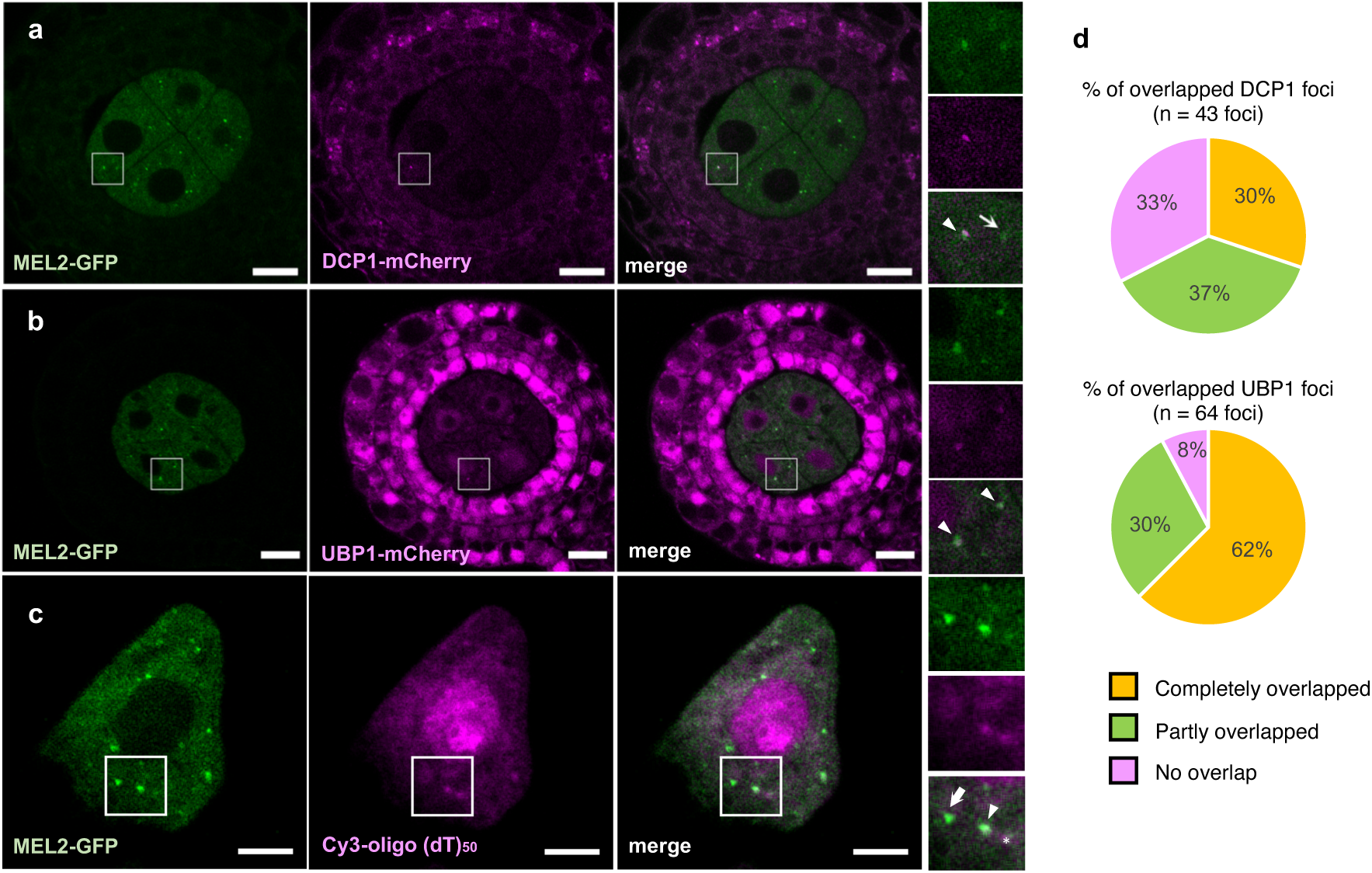
MEL2 granules are associated with P-bodies, stress granules, and poly(A)+ RNA in PMCs. (**a, b**) Representative confocal images of MEL2-GFP with DCP1-mCherry (**a**), UBP1b-mCherry (**b**) in cross-sections of 0.4-mm (PL stage) anther lobes in transgenic rice plants. (**c**) Representative confocal images of MEL2-GFP with Cy3-labeled poly(A)+ RNAs in PMC isolated from 0.4-mm (PL stage) anther. The rightmost three panels are magnified views of white squares in larger panels: from top to bottom, GFP, mCherry, and merged images. Arrows indicate sole MEL2 granules and arrowheads indicate MEL2 granules colocalized with DCP1, UBP1b, or poly(A)+ RNA. Bars = 10 μm in (**a**) and (**b**); 5 µm in panel (**c**). (**d**) Pie charts displaying the percentage of DCP1-mCherry or UBP1-mCherry foci overlapping with MEL2-GFP granular foci.

With OsUBP1b, both nuclear and cytoplasmic granule signals were more prominent than with OsDCP1 in both PMCs and somatic wall cells of PL anthers (Fig. 4b, Supplementary Fig. 5c). In PMCs, 62.5% and 30.0% of 64 cytoplasmic OsUBP1b foci completely and partially overlapped with MEL2 granular foci, respectively (Fig. 4b, d, Supplementary Fig. 5c). Notably, this strong colocalization was observed in plants grown normally in the absence of heat treatment.

These results suggest that MEL2 granules tend to act together with stress-granule components, rather than P-bodies, although it remains possible that MEL2 interacts in some fashion with P-bodies.

### MEL2 granules are associated with polyadenylated transcripts

Next, we investigated whether MEL2 granular foci contain polyadenylated RNAs. To this end, we performed fluorescence *in situ* hybridization (FISH) analysis using a Cy3-labeled poly(dT)_50_ nucleotide probe. In PMCs at the premeiotic stage, Cy3 signals were detected in both the nucleus and cytoplasm (Fig. 4c, Supplementary Fig. 5d). In the cytoplasm, several Cy3-dense signals were observed, a subset of which overlapped with MEL2 granules. By contrast, signals from Cy3-labeled poly(dA)_50_, used as a negative control, were barely detectable (Supplementary Fig. 5e). Thus, we concluded that the aggregated foci of MEL2-GFP are specific to the RNA granules spatiotemporally limited to the cytoplasm of premeiotic PMCs.

### Robust formation of MEL2 granules requires interactions of all conserved domains

In addition to ankyrin repeats, an RRM domain, and C3HC4-type RING domains [21], we found that MEL2 contains a conserved winged helix-turn-helix (wHTH)-containing LOTUS domain between the RRM and RING domains (Fig. 5a, Supplementary Fig. 7). The LOTUS domain is conserved in many factors associated with germ granules in animals (Supplementary Fig. 7) [28]. To clarify the function of the four MEL2 domains in granule formation, we fused each domain with GFP and overexpressed these constructs in rice protoplasts under the control of the constitutive *Actin* promoter (Fig. 5a). GFP fluorescence patterns in protoplasts were divided into four classes; the pattern of intact MEL2-GFP (FL) was defined as class I (Fig. 5a, b). ANK, RRM, or LOTUS alone failed to form cytoplasmic granules (class IV) (Fig. 5b, c), whereas RING alone exhibited granular structures accompanied by faint diffuse signals (class II) in 16 of 51 cells (Fig. 5c). Conversely, when a mutated MEL2 protein lacking any of the four conserved domains was introduced, all constructs, including ΔRING, formed granules, although the diffuse cytoplasmic signal was more intense in ΔRING than in the other three (Fig. 5d). Next, we introduced constructs consisting of each pair of consecutive domains: ANK–RRM (ARr), RRM–LOTUS (RrL), and LOTUS–RING (LRi). In all cases, we observed indistinct granular foci with moderate diffuse signals (class III) (Fig. 5b, e). The ARrL construct, in which LOTUS was added to ARr, formed more robust class II granules than class III granules (Fig. 5e), suggesting that LOTUS, in addition to RING, makes a significant contribution to MEL2 granule formation.

**Fig. 5.**
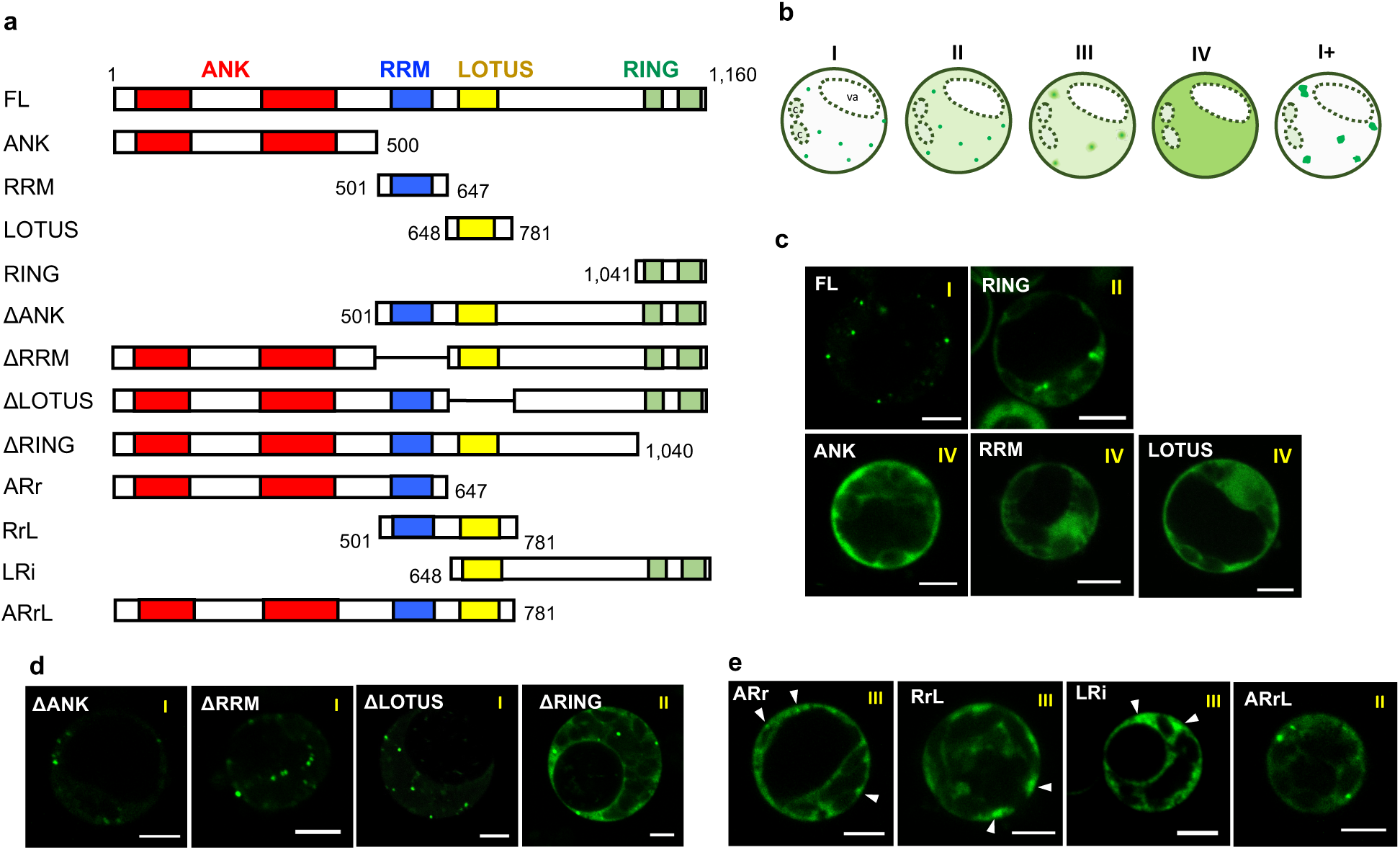
Truncated protein assay of MEL2 for granule formation in rice protoplast cells. (**a**) Schematic diagram of primary protein structures of full-length and truncated MEL2. (**b**) Schematic illustration of five patterns of MEL2-GFP subcellular localization: class I, robust granules only; II, granules, faint diffusive signals; III, indecisive granules with diffusive signals; IV, strong diffusive signals only; I+ (Fig 6), excess aggregation of granules. va, vacuole; c, chlorophyll. (**c–e**) Representative subcellular localizations of various truncated versions of MEL2 (**a**) fused with GFP in protoplasts. Arrowheads in (**e**) indicate class III indecisive granules. Bars = 5 μm.

### A spacer domain that includes part of the intrinsically disordered region regulates the stability of MEL2 granule

ΔANK, ΔRRM, ΔLOTUS, ΔRING, and LRi, all of which had the potential to form class II or III granules, shared a long domain-less spacer region between LOTUS and RING. This region contains nine <u>p</u>roline-<u>r</u>ich and tandemly <u>r</u>epeated peptides (PRRs), each consisting of 11 or 12 conserved amino acids, RKPx [V/L]IEPVPT (Supplementary Fig. 8a, b). To verify the importance of the spacer region in granule formation, we introduced a C-terminal spacer (SPC)-deleted MEL2 protein (ΔSPC) into protoplasts. Intriguingly, ΔSPC formed significantly enlarged granules (class I+) (Fig. 5b, 6e, Supplementary Fig. 8c, d), implying a suppressive role for the SPC or PRRs in MEL2 aggregation.

**Fig. 6.**
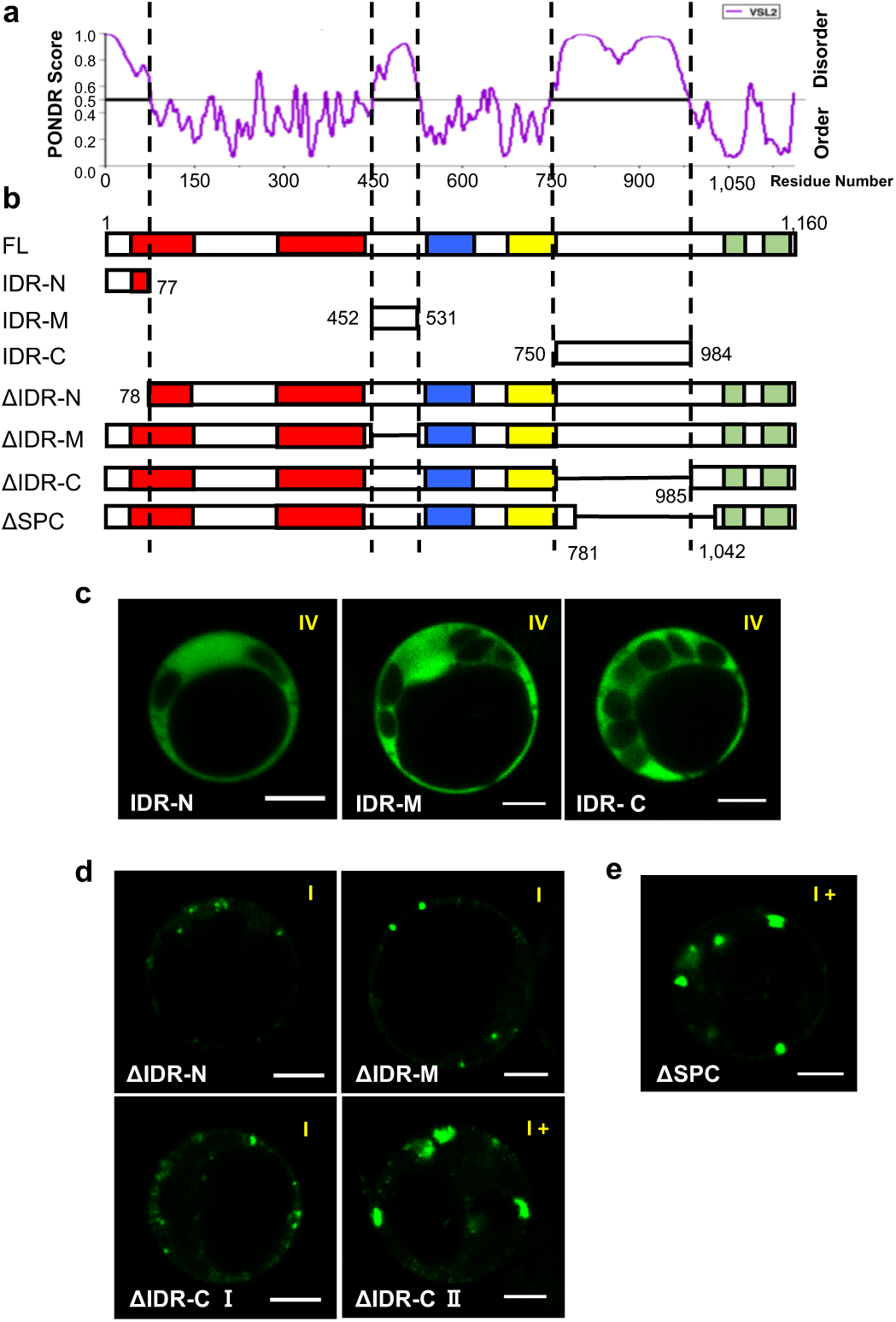
The impact of three IDRs and the spacer region on MEL2 granular size. (**a**) Prediction of intrinsically disordered regions (IDRs) in MEL2 protein by the PONDR-VSL2 algorithm. (**b**) Schematic diagrams of truncated MEL2 proteins in (**c–e**). (**c–e**) Representative images of truncated MEL2-GFP fusion proteins in protoplasts. Bars = 5 μm.

In general, the RNP granule behaves like a liquid droplet assembled by means of liquid–liquid phase separation (LLPS), the mechanism underlying the formation of membrane-less organelles [29, 30]. The intrinsically disordered region (IDR) lacks a stable folded structure and engages in weak multivalent protein–protein and protein–RNA interactions that contribute to LLPS [31, 32]. When the MEL2 peptide sequence was parsed with PONDR-VSL2, a prediction algorithm for IDRs [33], three putative IDRs were found: IDR-N, IDR-M, and IDR-C (Fig. 6a, b). The IDR-C region spanned residues 782–984 within the SPC that contains the nine PRRs (Supplementary Fig. 8a). We transcriptionally fused each of the three IDRs with GFP and expressed them in protoplasts, but all yielded class-IV diffuse signals in the cytoplasm (Fig. 6c), suggesting that each individual IDR has little granule formation potential. Next, we removed individual IDRs from MEL2 protein and introduced the resultant constructs into protoplasts. ΔIDR-N and ΔIDR-M formed class-I granules, comparable to those formed by wild-type MEL2 (Fig. 6d). ΔIDR-C also formed granules, but their sizes fluctuated and appeared to be separable into two types. Most cells contained class-I granules (Fig. 6d, bottom left), whereas 9.8% of cells (4/41) contained class-I+ granules (Fig. 6d, bottom right), similar to the results obtained with ΔSPC (Supplementary Fig. 8c, d).

Given that ΔSPC formed stabler class-I+ granules than ΔIDR-C, we concluded that the C-terminal end of SPC (a.a. 986–1042), rather than the PRRs, plays a predominant role in suppressing excess aggregation to maintain granule stability.

### MEL2-like proteins with diversified domain combinations are conserved widely in the plant kingdom

Next, we searched for MEL2-like (MEL2L) proteins in other plant species using BLASTP with full-length MEL2 as the query sequence. The phylogenetic relationships of MEL2 and MEL2L proteins, with their summarized domain compositions, are shown in Fig. 7A. The analysis revealed that MEL2L proteins with varied domain composition are conserved in many land plants and charophyte alga *Chara braunii,* the extant algal species phylogenetically closest to land plants. No MEL2L protein was detected in the outgroup chlorophyte *Chlamydomonas reinhardtii,* indicating that MEL2L emerged in charophytes after their divergence from chlorophytes. This analysis revealed another conserved domain, a C3H1-type zinc finger motif (ZINC), between ANK and RRM of MEL2Ls (Fig. 7a). In light of recent studies that highlighted the role of ZINC in RNA binding, in addition to DNA binding [34], this domain may participate in RNA binding alongside RRM. If ZINC is considered the fifth domain, the consensus domain composition of MEL2L can be defined as ANK–ZINC–RRM–LOTUS–RING.

**Fig. 7.**
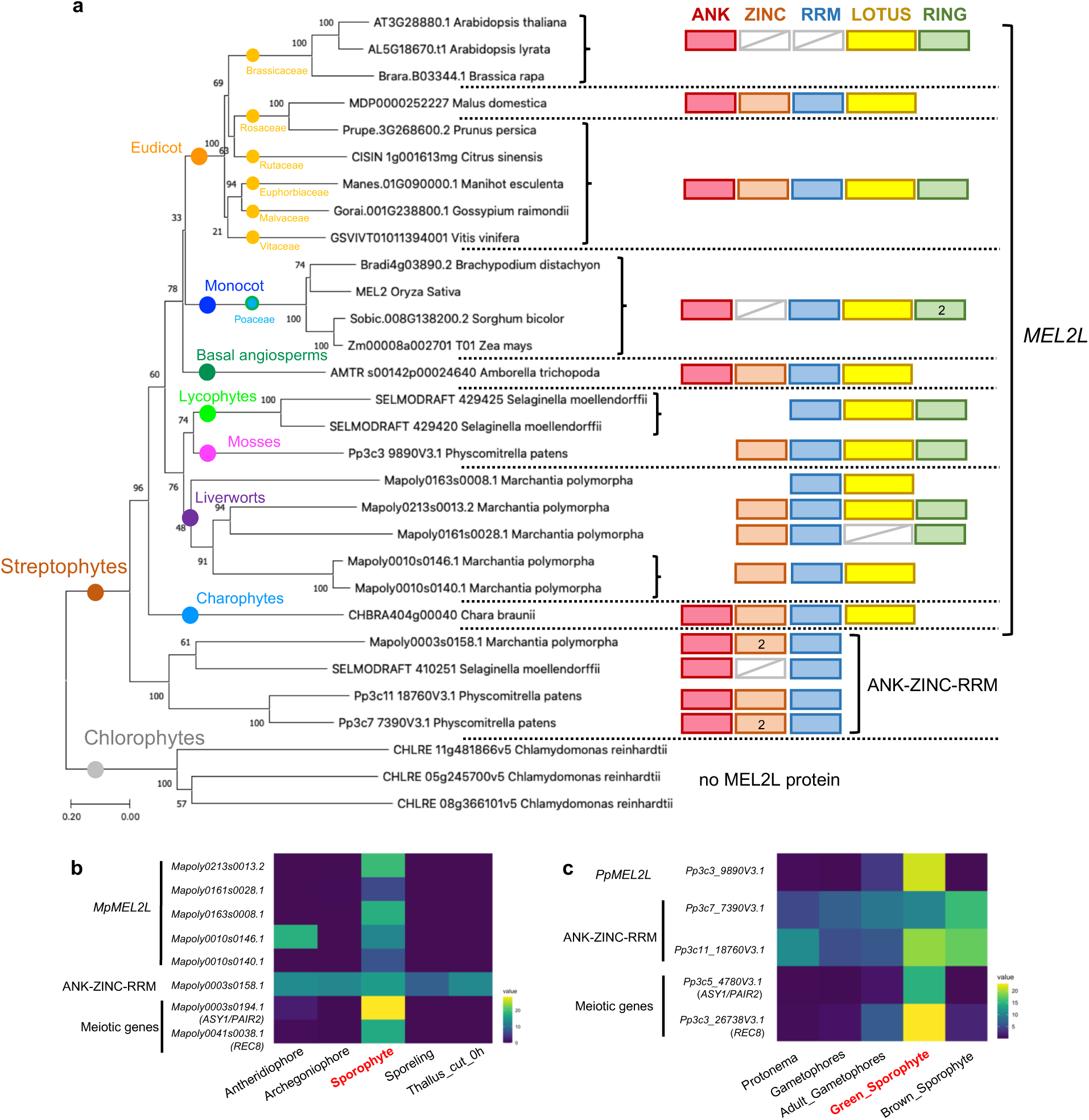
MEL2-like proteins are widely conserved in the plant kingdom. (**a**) Phylogenetic tree of MEL2-like proteins with simple domain architecture widely conserved in the plant kingdom. (b) Expression profile of *Marchantia polymorpha* MEL2-like genes in various tissues. FPKM values of RNA-seq analysis from Bowman et al. (2017) were used to construct the heatmap [35]. *Mapoly0003s0194* and *Mapoly0041s0038* encoding ASY1/PAIR2 and REC8 orthologs, respectively, are shown together as meiotic markers. (c) Expression profile of *Physcomitrella patens* MEL2-like genes in various tissues. Expression values were obtained from the PEATmoss website (https://peatmoss.online.uni-marburg.de/index) and used to construct the heatmap [36]. *Pp3c5_4780V3.1* and *Pp3c3_26738V3.1*, encoding *ASY1/PAIR2* and a *REC8* ortholog, respectively, are shown as meiotic markers. Sporophyte tissues presumed to be at the meiotic stage are described in red letters in (**b**) and (**c**).

The domain composition of rice MEL2, ANK–RRM–LOTUS–RING, is conserved only in Poaceae species (Fig. 7a). A domain combination consistent with the MEL2L consensus was found in Rosaceae, Rutaceae, Euphorbiaceae, Malvaceae, and Vitaceae. The charophyte *Chara braunii*, the basal angiosperm *Amborella trichopoda*, and the eudicot *Malus domestica* contained RING-lacking MEL2Ls (ANK–ZINC–RRM–LOTUS). Interestingly, basal land plants (*M. polymorpha* and *Physcomitrella patens*) and *S. moellendorffii* contained multiple MEL2Ls lacking ANK or RING (Fig. 7a). The MEL2Ls of Brassicaceae uniquely lacked both RRM and ZINC. All MEL2L proteins possessed the RRM and LOTUS domains, with the exception of three Brassicaceae MEL2Ls and one of five *M. polymorpha* MEL2Ls (Mapoly0161s0028.1), suggesting that the RRM–LOTUS pair plays core roles in the molecular function of MEL2Ls. The five *M. polymorpha* genes and one *P. patens MEL2L* gene were expressed exclusively at the sporophyte, in which genes orthologous to *Arabidopsis* and rice meiotic genes (*ASY1/PAIR2*, *REC8*) were abundantly expressed (Fig. 7b, c) [35, 36]. These results indicate that the expression of MEL2Ls containing RRM and/or LOTUS is regulated strictly and spatiotemporally in basal land plants.

In addition to MEL2Ls, we found that *M. polymorpha*, *P. patens*, and *S. moellendorffii* also retained ANK–ZINC–RRM-encoding and LOTUS-lacking proteins, *Mapoly0003s0158.1*, *Pp3c7_7390V3.1*, *Pp3c11_18760V3.1* and *SELMODRAFT410251* genes (Fig. 7a). However, *M. polymorpha* and *P. patens* genes of them were excluded from the MEL2L family because they were phylogenetically grouped outside the MEL2L clade (Fig. 7a), consistent with their ubiquitous expression throughout the lifecycle (Fig. 7b, c) [35, 36].

Curiously, the BLASTP search failed to detect MEL2L proteins in Fabaceae and Solanaceae. Furthermore, *Arabidopsis* and *Brassica rapa* contained MEL2Ls lacking both ZINC and RRM, in contrast to other species (Fig. 7a). In rice, for example, *MEL2* expression was limited in reproductive organs (Supplementary Fig. 9) [37]. However, the putative *Arabidopsis MEL2L* gene, *At3g28880*, was expressed ubiquitously, with the highest expression levels in roots and internodes (Supplementary Fig. 10) [38].

Collectively, these observations show that MEL2L proteins are conserved in many extant species of land plants and charophyte algae, and that their biological functions are likely coupled with sporophytic developmental processes, including meiosis. However, Brassicaceae, Fabaceae, and Solanaceae species were exceptional in regard to the domain conservation or transcriptional patterns of MEL2L proteins and genes, despite the fact that all land plants share a sporophytic phase and meiosis.

## Discussion

In this study, we found that MEL2-GFP formed aggregates in the cytoplasm of premeiotic PMCs in transgenic rice plants (Fig. 2–4) and ectopically in rice leaf protoplasts (Supplementary Fig. 4). MEL2 granules associated with poly(A)+ RNAs at the premeiotic PMC cytoplasm *in vivo* (Fig. 4). These findings indicate that rice MEL2 is a component of cytoplasmic RNP granules, and that its expression is spatiotemporally and strictly regulated to ensure proper meiotic entry of rice germ cells. It remains possible that MEL2 diffusing in the intergranular cytoplasmic space has other functions.

The germ granule is a general term for a cytoplasmic, non– membrane-bound, and RNA-rich organelle that functions in the control of germ cell specification, formation, and migration [39, 40]. In *Drosophila* and *C. elegans*, germ cell specification to establish the germline is achieved by germ granules in early embryos. By contrast, in mammals it occurs long after embryogenesis, and germ granules function in germ cell differentiation, rather than specification [39]. Like mammals, most land plants develop germ cells at late stages. Thus, the rice MEL2 granule might be comparable to the mammalian germ granule, which is required for differentiation and meiotic transition of spore mother cells.

The molecular function of germ granules and their components has been implicated in translational control and RNA stabilization or degradation [39]. Intriguingly, if the information from all plant genomes was removed, a TBLASTN search using the MEL2 RRM as a query returned human DAZ-associating protein 1 (DAZAP1) as highly similar (AF181719, Score; 57 bits, E-value; 2e-7) [21]. Although DAZAP1 has not been reported to be a germ granule component, it is highly expressed in the testis and stimulates the translation of target mRNA [41]. DAZAP1 is potentially associated with stress granules in puromycin-treated iPS cells [42]. Furthermore, when the RRMs of DAZAP1 are swapped into G3BP1, a core factor of the stress granule, the resultant chimeric protein is functional in stress granule assembly [43]. DAZL, a potential binding partner of DAZAP1, is an RRM-containing protein widely conserved in vertebrates [25, 44], which promotes meiosis by activating the expression of the meiotic inducer Stra8 in mouse [45]. DAZL localizes to stress granules as well as DAZAP1 by heat treatment, which are hypothesized to protect spermatogonia and early spermatocytes from heat stress by inhibiting apoptotic signaling and translational repression [46]. The association of rice MEL2 with stress granules is likely analogous to that of the RRM-containing mammalian proteins DAZAP1 and DAZL, which have been implicated in translational control of target RNAs. Therefore, it is plausible that MEL2 granules capture and store meiotic target mRNAs to temporarily repress their translation until meiotic entry. However, it is also possible that MEL2, like DAZL, stimulates the translation of meiotic genes [47, 48].

Recent works showed that the gene expression profile changes dramatically at the early meiotic prophase I in maize PMCs [49], and the rice reproductive phased siRNAs (phasiRNAs) associated with MEL1 Argonaute regulate the reprogramming of gene expression through the direct RNA cleavage [50, 51]. In addition to phasiRNAs-MEL1 mediated mRNA degradation, some MEL2 granules may be involved in the decay process working alongside P-bodies to remove a subset of transcripts that are dispensable for progression through meiosis.

We found that MEL2 contains three IDRs in addition to four functional domains (Fig. 5 and 6). IDR-C is the largest of three MEL2 IDRs, and we speculated that it has a repressive function in MEL2 granular size. Distinct IDRs with various physiological properties are involved in phase separation in different manners [30–32]. Given that the nature of an IDR is an important determinant of the state of granules (i.e., liquid-like or solid-like) and dynamic behaviors such as fission and fusion, it is possible that MEL2 ΔIDR-C and ΔSPC may alter the MEL2 granule in favor of the solid state, resulting in excess aggregation due to a conformational change or misfolding of the MEL2 protein. Alternatively, these regions may be responsible for interactions with RNAs or proteins to modulate physiological states and prevent the granule from engaging in undesirable interactions or excess protein/RNA aggregation. Further *in vivo* and *in vitro* analyses will be necessary to elucidate the function of the largest MEL2 IDR and its involvement in intra- and intermolecular interactions.

Our analysis revealed that MEL2-like genes are widely conserved in land plants and charophyte algae with diverse domain structures. Green plants comprise two lineages, the streptophytes that colonized land and the chlorophytes that have adaptations for land but remained mostly aquatic [52]. All land plants are thought to have evolved from aquatic charophyte algae, implying that the prototype MEL2 evolved after the divergence from chlorophytes. The function of MEL2L in charophytes is unknown, but it may be involved in temporal arrest of zygotic meiosis until the environment is suitable for meiosis and subsequent germination. One possibility is that meiotic arrest or delay caused by MEL2L protein may allow intercalation of mitosis before meiosis, leading to establishment of the sporophytic phase in land plants, although verification of this attractive idea would require additional evidence.

From phylogenetic analysis, the consensus composition of MEL2L domains could be defined as ANK–ZINC–RRM–LOTUS–RING, and a consecutive RRM–LOTUS pair is conserved in all land plants and *C. braunii*, with the exception of Brassicaceae (Fig. 7a). Of the deleted versions of MEL2 that we examined in this study, RRM–LOTUS (RrL) had the ability to form class II granules (Fig. 5). These results imply that RRM–LOTUS constitutes a core that specifies MEL2 function in premeiotic spore mother cells. In turn, this means that other MEL2L domains do not always need to be packaged within a single protein with the RRM–LOTUS pair. MEL2L proteins may have evolved while repeating the domain shuffling of MEL2 domains other than the RRM–LOTUS core, and in some plant species, MEL2 function may have been assigned to multiple proteins that share these domains. The MEL2 domain shuffling seems to have occurred discontinuously in phylogenic lineages, as no MEL2L protein was found in Fabaceae and Solanaceae, both of which are eudicot species, or in Rutaceae and Vitaceae, which conserve MEL2L proteins (Fig. 7a).

Domain shuffling, which is thought to be driven by exon shuffling in eukaryotes, has been identified as one of the major mechanisms for creating new proteins by rewiring pre-existing domains over the course of evolution (reviewed in Souza 2012) [53]. The current model has proposed that a “promiscuous” domain, meaning a hub for physical protein–protein interactions (PPIs), tends to be reused repeatedly in exon or domain shuffling in complex PPI networks, although this trend is observed more often in metazoan species, and on a much smaller scale in *Arabidopsis* [54]. The fact that MEL2 domains contribute almost equally to MEL2 granule formation (Fig. 5) is likely advantageous in the shuffling process; even if MEL2 functions and domains were shared with multiple proteins, and several of these proteins were to vanish during evolutionary processes, MEL2 granules could be reconstructed by gathering together multiple proteins that share the necessary domains. Indeed, the MEL2-like *P. patens* (Pp3c3_9890V3) and *Arabidopsis* (At3g28880) proteins have architectures similar to those of MEL2 ΔANK and ΔRRM, respectively (Fig. 5 and 7a). Given that both ΔANK and ΔRRM could form cytoplasmic granules in rice protoplast cells, we predict that these proteins have the potential to form granules.

In summary, we propose that the MEL2 granule modulates the cell cycle to ensure proper meiotic transition, based on its ability to associate with poly(A)+ RNAs and components of the P-body and stress granules. Colocalization of MEL2 with UBP1b, a stress-granule component, suggests that MEL2 granule formation may be stimulated by intrinsic signals such as reactive oxygen species, the levels of which are elevated in meiotic anthers in rice [55]. In addition to granule induction, the unknown signaling factors that cause timely and rapid destruction of MEL2 granules in PMCs are also of interest in considering the mechanism of meiotic entry. Further investigations will illuminate the importance of RNA granules, including MEL2 granules, in plant reproduction, and reveal similarities and differences in the properties of premeiotic and meiotic germ cells between animals and plants.

## Methods

### Plant materials

Rice plants carrying the *mel2-1* mutant allele, which harbors an insertion of the endogenous *Tos17* retrotransposon [21], were backcrossed five times with cv. Nipponbare. Genotypes of plants segregating in BC5F2 populations were determined by PCR as described by Nonomura et al (2011) [21]. Wild-type cv. Nipponbare and *mel2-1* plants were grown in open paddy fields at the National Institute of Genetics (NIG), Mishima, Japan. Transgenic plants were grown in a growth chamber at 30°C for 14 hr with light and at 25° for 10 hr in the dark.

For protoplast experiments, Nipponbare seedlings were grown on 1/2 MS medium for 7–9 days in a growth chamber at 28°C for 14 hr with light and at 25° for 10 hr in the dark.

### Construction of transgenic plants expressing fluorescently tagged protein

For genomic *MEL2-GFP* fusion, genomic fragments of 1.5 kb upstream and 1.7 kb downstream from the *MEL2* stop codon were amplified by PCR using the Nipponbare genome as a template and gene-specific primers, including the restriction enzyme sites. They were inserted into the CaMV35S-sGFP(S65T)-nos3’ vector [56] at the *Hind*III/*Nco*I and *Bsr*GI/*Eco*RI sites such that the *GFP* CDS was fused in-frame to the C-terminal *MEL2*-coding sequence. The resultant 3.9-kb genomic *MEL2-GFP*–fused sequence was excised by digestion with *Age*I and *Xba*I. Next, the 7.1-kb fragment containing the *MEL2* promoter, the 5’ UTR, and the *MEL2*-coding sequence was excised by digestion with *Sal*I and *Age*I from the genomic *MEL2*-containing plasmid used for complementation of the *mel2* sterile phenotype in a previous study [21]. The 7.1-kb and 3.9-kb fragments were inserted together into pPZP2H-lac binary vector at the *Sal*I/*Xba*I sites, as the *MEL2* terminus was replaced with the in-frame *MEL2-GFP* sequence. The MEL2-GFP construct was transformed into the *mel2-1* background as described in Nonomura et al., (2011) [21].

*OsDCP1* (*Os12g0156400*) and *OsUBP1b* (*Os11g0620100*) genes were identified on the rice genome by BLASTp search on the RAP-DB website (https://rapdb.dna.affrc.go.jp) using the *Arabidopsis DCP1* and *UBP1b* genes as query sequences [57, 58]. The 7.0-kb and 9.0-kb genomic fragments of *OsDCP1* and *OsUBP1b* (including 1.5-kb and 2.2-kb sequences upstream of the transcriptional start sites [TSS]) were PCR-amplified and sub-cloned into pBluescript II SK(+), respectively, and the PCR-amplified *mCherry* sequence was inserted in frame before the stop codon of both rice genes using NEBuilder HiFi DNA Assembly Master Mix (New England Biolabs). The genomic *OsDCP1-mCherry* and *OsUBP1b-mCherry* sequences were cut out by digestion with *Kpn*I/*Xho*I and *Kpn*I/*Not*I, respectively, transferred into binary vector pRI909 (Takara Bio), transformed into *Agrobacterium* strain EHA105, and used for transformation of plants retaining the MEL2-GFP transgene. The transformants were selected with the antibiotic G418. Primers used for these constructions are listed in Supplementary Table 2.

### Construction of deletion series and transient expression analysis of rice protoplast cells

Total RNA was extracted from 0.2–0.5-mm anthers, and a cDNA library was created using the SuperScript III First-Strand Synthesis System (Thermo Fisher Scientific). The MEL2 protein–coding sequence (CDS) without a stop codon was PCR-amplified with primers in which the *Apa*I and *Xba*I sites were added to the 5’- and 3’-ends of MEL2 CDS, respectively. The amplicon was cloned into the pCR-Blunt II TOPO vector (Invitrogen, MEL2-FL-TOPO). Next, the *OsACTIN* promoter (*OsACT*p) [59] was PCR-amplified and cloned into the *GFP*-containing pUC119 vector, as *Apa*I and *Xba*I sites were present between *OsACT*p and *GFP* (pACTp-A/X-GFP). After *Apa*I/*Xba*I digestion of MEL2-FL-TOPO, the MEL2 CDS fragment was inserted into pACTp-A/X-GFP, as *GFP* was fused in frame with *MEL2*. For truncation analysis, partial MEL2 sequences were amplified with primers containing *Apa*I or *Xba*I sites using MEL2-FL-TOPO as the template. The amplicons were digested with *Apa*I and *Xba*I and inserted in-frame between the Actin and GFP sequences. To delete a single domain, inverse PCR was carried out using MEL2-FL-TOPO as a template, and the amplicons were self-ligated. The domain-deleted MEL2 CDS were cut out and inserted as described above.

For construction of mCherry-tagged protein, the Actin promoter was PCR-amplified and cloned into mCherry containing pBluescript SK(+) vector such that *Kpn*I/*Spe*I sites were present between the two sequences (pACTp-K/S-mCherry). Using the anther cDNA library, *OsDCP1* and *OsUBP1b* CDS were amplified using primers that added *Kpn*I and *Spe*I sites to the 5’-end and 3’-end of the respective CDSs without stop codons, and then cloned into the pCR-Blunt II TOPO vector (*OsDCP1*-TOPO, *OsUBP1b*-TOPO). After *Kpn*I/*Spe*I digestion of *OsDCP1*-TOPO and *OsUBP1b*-TOPO, the *OsDCP1* and *OsUBP1b* CDS fragments were inserted individually into pACTp-K/S-mCherry. The primers used for these constructions are listed in Supplementary Table 2.

The constructs described above were co-transfected into protoplast cells prepared from young leaves of rice seedlings. Transfections were performed by the PEG method, as described by Zhang et al. (2011) [60].

### Observation of fluorescently tagged proteins

Young rice panicles, including premeiotic and meiotic flowers, were placed into a 50 mL screw-cap tube and fixed with 4% paraformaldehyde/PBS (pH 7.4) (4%PFA) four times for 20 min each under vacuum on ice. After fresh fixative was added, the tube was gently shaken for 2 hr. The fixed panicles were washed with PBS six times for 20 min each and kept at 4°C until use. After removal of the palea and lemma, the floret with exposed anthers was embedded in 6% agarose gel and sliced to a 50-µm thickness on a MicroSlicer DTK-ZERO1 (D.S.K.). Fluorescence signals were visualized on a Fluoview FV300 CLSM system (Olympus), as described by Ono and Nonomura (2018) [61].

### Indirect immunolocalization and image analysis

Indirect immunofluorescence staining was performed as described previously [27] with minor modifications. Briefly, to observe PAIR2 signals, MEL2-GFP– expressing PMCs were isolated from premeiotic and meiotic anthers fixed with 4% PFA and incubated with mouse anti-PAIR2 antibody (1:3000) overnight at 4°C in a dark box. The suspension was replaced with Cy3-conjugated anti–mouse IgG (1:200) and incubated for 3 hr at room temperature. Signals were captured on a Fluoview FV300 CLSM system (Olympus). The number and size of MEL2 granules were measured using binarized captured images and Particle Analyzer implemented in the Fiji platform [62]. The intensity of the Cy3-stained PAIR2 signal in each nucleus was measured in the Fiji platform, and normalized against nuclear area.

### Fluorescence *in situ* hybridization

RNA FISH analysis was performed as described by Hyjek-Składanowska et al. (2020) [63] with minor modifications. Anthers containing PMCs expressing MEL2-GFP, fixed in 4% PFA/PBS, were incubated for 10 min in the same enzyme cocktail used for indirect immunolocalization, and then squashed on a glass slide to release PMCs. The enzyme cocktail contained 2% cellulase Onozuka-RS (Yakult), 0.3% Pectolyase Y-23 (Kikkoman), 0.5% Macerozyme-R10 (Yakult), and 3.75 µg/µl Cytohelicase (Sigma-Aldrich). The cells were permeabilized with 0.1% Triton X-100 in PBS for 20 min, followed by three washes with 1× PBS. Hybridization buffer (30% v/v formamide, 4× SSC, 5× Denhardt’s solution, 1 mM EDTA, and 50 mM phosphate buffer) containing 50 pmol/ml Cy3-labeled oligo probes was applied to the cells and incubated overnight at room temperature. The cells were then washed five times with 4× SSC and five times with 2× SSC / 1× PBS, each for 1min. Signals were captured on a Fluoview FV300 CLSM system (Olympus).

### mRNA-seq analysis

For mRNA-seq analysis, anthers at the PE, PL, and ME stages were collected from wild-type and *mel2* mutant flowers, immediately frozen with liquid nitrogen in microtubes, and stored at −80°C. Three biological replicates were prepared for every stage of every plant type, and approximately 180 anthers were used for each replicate. Total RNA was extracted using the NucleoSpin RNA Plant (Takara Bio) kit. One microgram of each RNA sample was used for library construction using the KAPA mRNA HyperPrep Kit (KAPA Biosystems). Eighteen libraries differentially indexed using the FastGene Adaptor kit (Nippon Genetics) were multiplexed in one lane and sequenced (2 × 150-bp paired-end reads) on a HiSeq X (Illumina) by GENEWIZ.

Adaptor and polyA tail sequences were removed from raw reads *in silico* using the Trimmomatic and PRINSEQ software [64, 65]. The resultant clean reads were mapped to the rice genome IRGSP-1.0 using HISAT2 [66]. The number of reads mapped to annotated genes was counted using featureCounts [67]. Differential expression analysis was carried out using DESeq2 [68]. Whole count data normalized by variance-stabilizing transformation (VST) are shown in Supplementary Table 3; 11 meiotic genes and *MEL2* were extracted for construction of heatmap. Differentially expressed genes were subjected to GO analysis using PANTHER at the Gene Ontology resource website (http://geneontology.org).

### Phylogenetic analysis

Amino acid sequences of MEL2-like proteins were obtained from Phytozome v12.1 (https://phytozome.jgi.doe.gov/pz/portal.html) or ORCAE (https://bioinformatics.psb.ugent.be/orcae/) by BLASTP search using MEL2 full-length peptide sequence as a query. Whole peptide sequences of each species were used for multiple alignments using MUSCLE embedded in the MEGA X software [69]. The phylogenetic tree was subsequently constructed by the neighbor-joining method with a bootstrap value of 1000. Three ankyrin repeat–containing proteins in *Chlamydomonas reinhardtii* identified by BLASTP search with the MEL2 full-length peptide sequence were used as an outgroup to place the root of the tree.

### Analysis of expression profile of MEL2 and MEL2L genes

mRNA-seq or microarray datasets for rice (*O. sativa*), *A. thaliana*, *M. polymorpha,* and *P. patens* were downloaded from Gene Expression Omnibus (GEO accession: GSE21396), the TraVA website (http://travadb.org), Bowman et al. (2017), and the PEATmoss website (https://peatmoss.online.uni-marburg.de/index), respectively [35–38]. Heatmaps were drawn using the ggplot2 package in R.

### Data availability

The data supporting the findings of this study are available within the paper and Supplementary Information files. mRNA-seq data in this study were deposited in the Sequence Read Archive of the DNA Data Bank of Japan (DDBJ) (accession number DRA011042). Identifiers of sequence data with sample descriptions are listed in Supplementary Table 4.

## Acknowledgements

We thank A. Otake, M. Kashihara and T. Hara for help with experimental work, field management and growing plants. We appreciate the National Bioresource Project (NBRP) Rice, conducted by the Japan Agency for Medical Research and Development (AMED) for providing us *mel2* mutant. This research was supported by MEXT KAKENHI (19H04871) and JSPS KAKENHI (18H02181) to K-I.N, and JSPS KAKENHI Grant-in-Aid JSPS Fellows (19J01894) and an NIG postdoctoral fellowship to M.M.

## Author contributions

M.M. and K-I.N. designed the research. M.M. performed almost all experiments. S.O. constructed the MEL2-GFP transgenic plants. M.M and K-I.N. analyzed the data and wrote the manuscript.

---

**Supplementary Figure 1.**
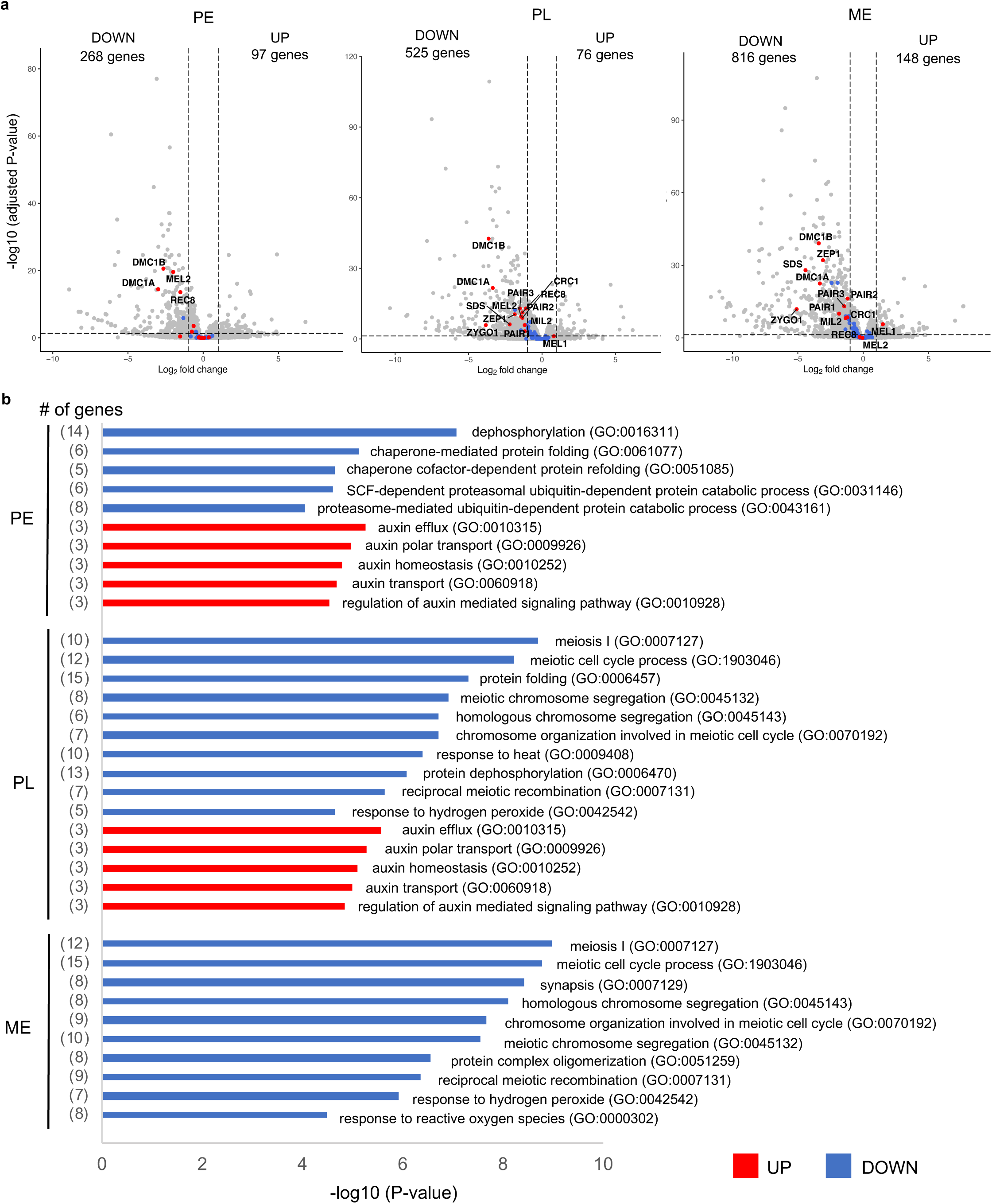
MEL2 is required for proper expression levels of meiotic genes. (**a**) Volcano plot displaying genes differentially expressed between wild-type and *mel2* mutant anthers at PL stage. The plot displays −log_10_(adjusted P-value) (y-axis) versus log_2_(fold change) (x-axis) of the transcript levels of 31283 coding genes. Horizontal and vertical dashed lines indicate the marginal lines of the cutoffs for adjusted P-value (0.05) and log_2_ fold change (−1 or 1). Gray spots represent all transcripts detected by mRNA-seq analysis. Blue spots represent genes with “meiotic cell cycle process” ontology (GO:1903046). Of these, 13 genes are labeled with red spots. (**b**) Gene Ontology (GO) enrichment analysis of genes differentially expressed in *mel2* mutant anthers. The data represented the −log_10_(P-value) of GO terms of biological processes found to be enriched in the gene set (Fisher’s exact test). GO terms and the number of genes are shown to the right and left of the bar graphs, respectively.

**Supplementary Figure 2.**
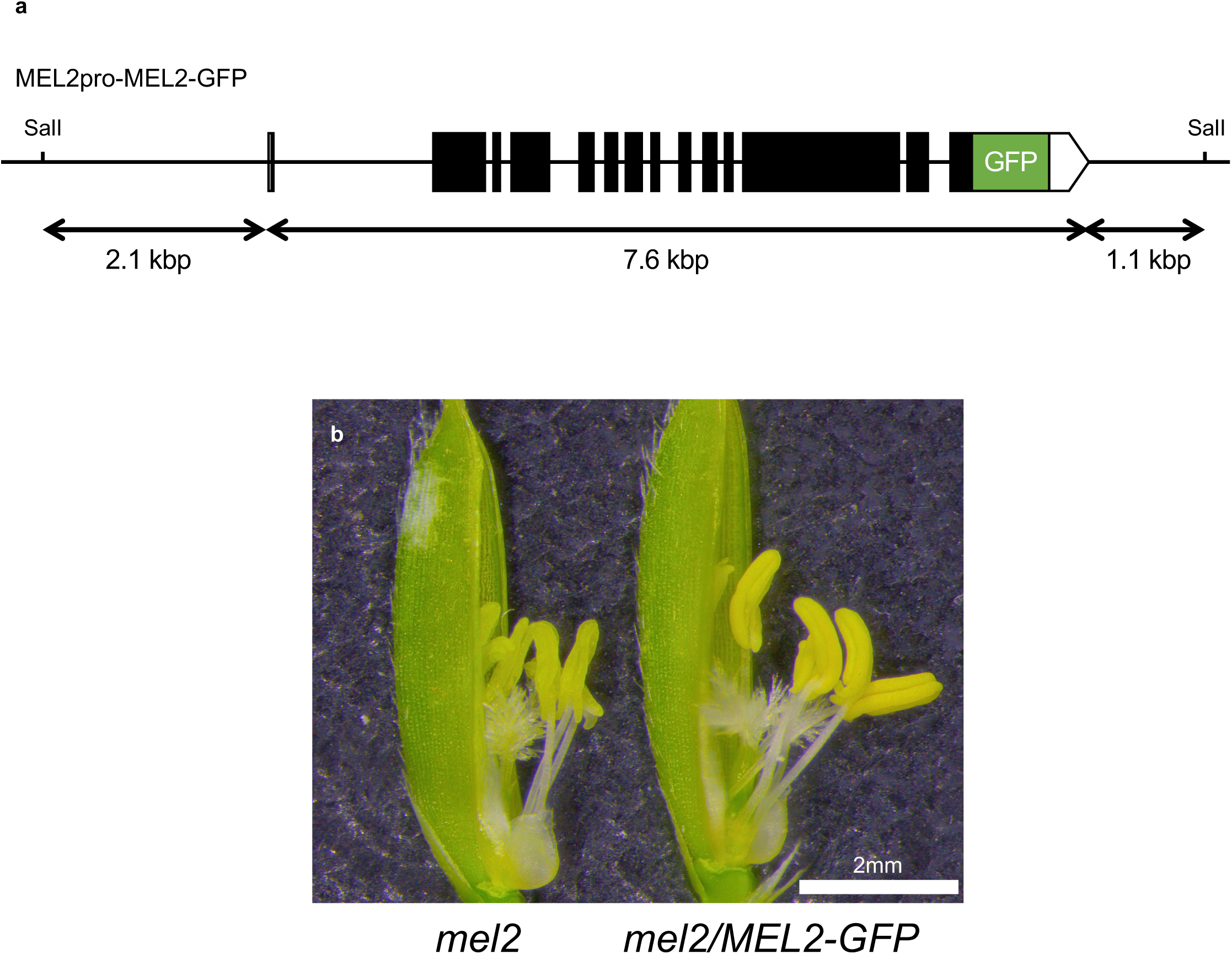
MEL2-GFP rescue *mel2* phenotype. (**a**) Diagram of proMEL2-MEL2-GFP transcriptional fusion construct. White boxes, black boxes, and green box represent UTRs, exons, and GFP, respectively. (**b**) *mel2* mutant and *mel2* mutant carrying the MEL2-GFP fusion construct. *mel2* anthers were shrunken and pale yellow, whereas MEL2-GFP transgenic plants had normal swollen anthers. Bar = 2 mm.

**Supplementary Figure 3.**
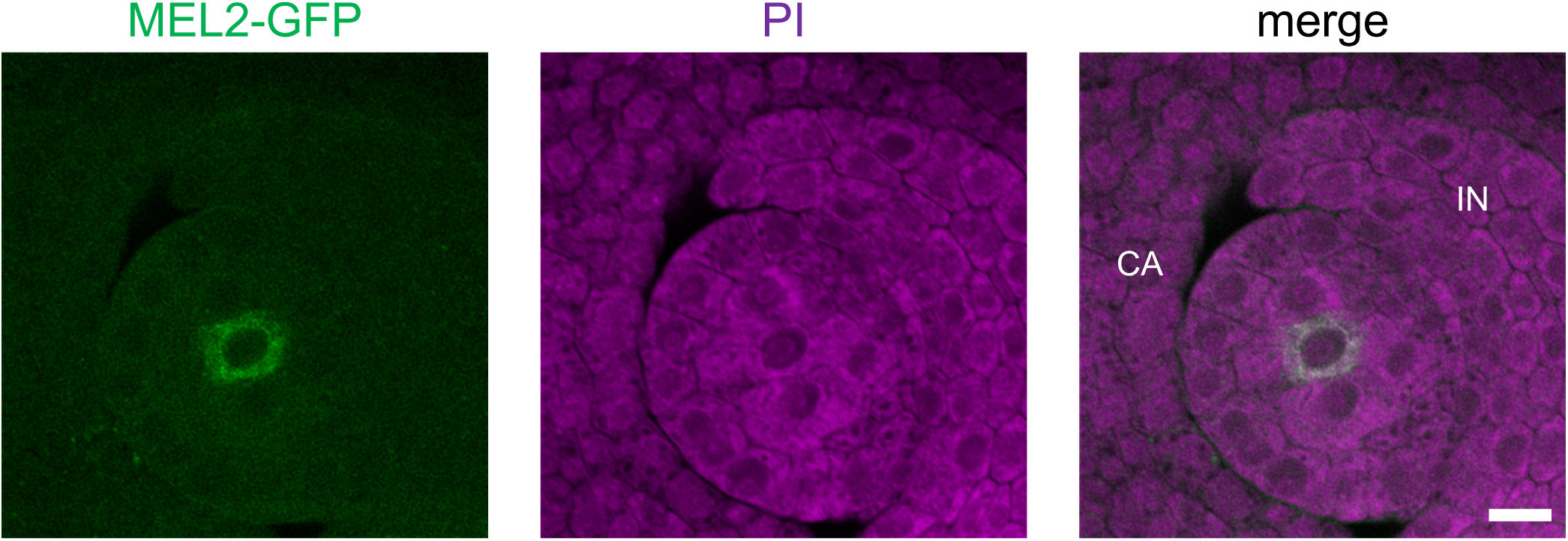
MEL2-GFP expression in female sporogenous cell. MEL2GFP signal was detected in the cytoplasm of the sporogenous cell at the center of the ovule, whereas little signal was observed in the surrounding cells. CA: carpel, IN: integument. Bar = 10 μm.

**Supplementary Figure 4.**
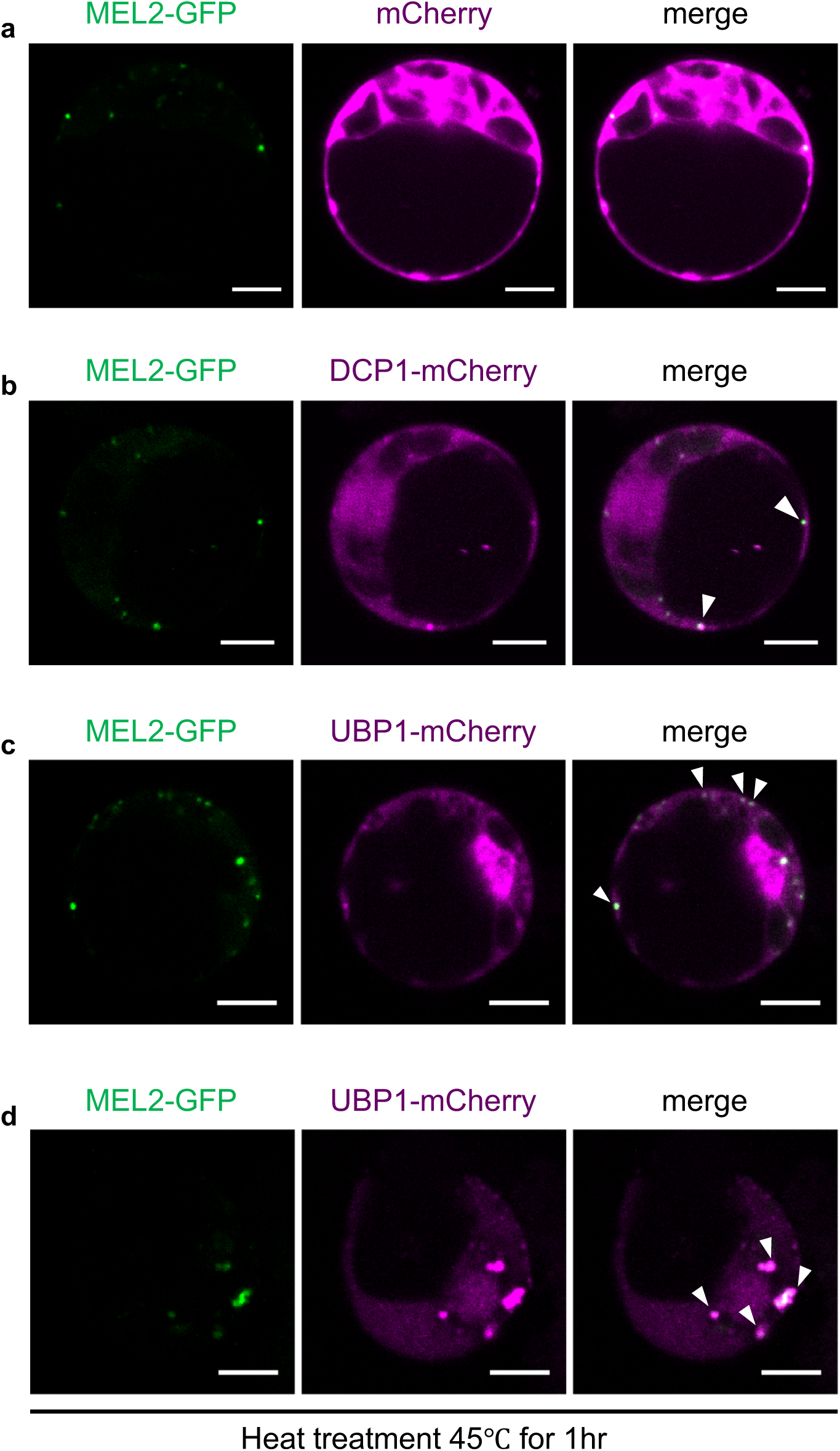
MEL2 granules associate with P-bodies and stress granules in rice protoplast cells. MEL2-GFP was co-expressed with free mCherry (**a**), DCP1-mCherry (**b**), and UBP1-mCherry (**c**, **d**) in rice mesophyll protoplast cells isolated from young seedlings (7–9 days old). All reporter proteins were driven by the constitutively expressed actin promoter. For (**d**), protoplast cells were subjected to heat stress (45°C) for 1 hr. White arrowheads indicate over-lapping granules. Bars = 5 μm.

**Supplementary Figure 5.**
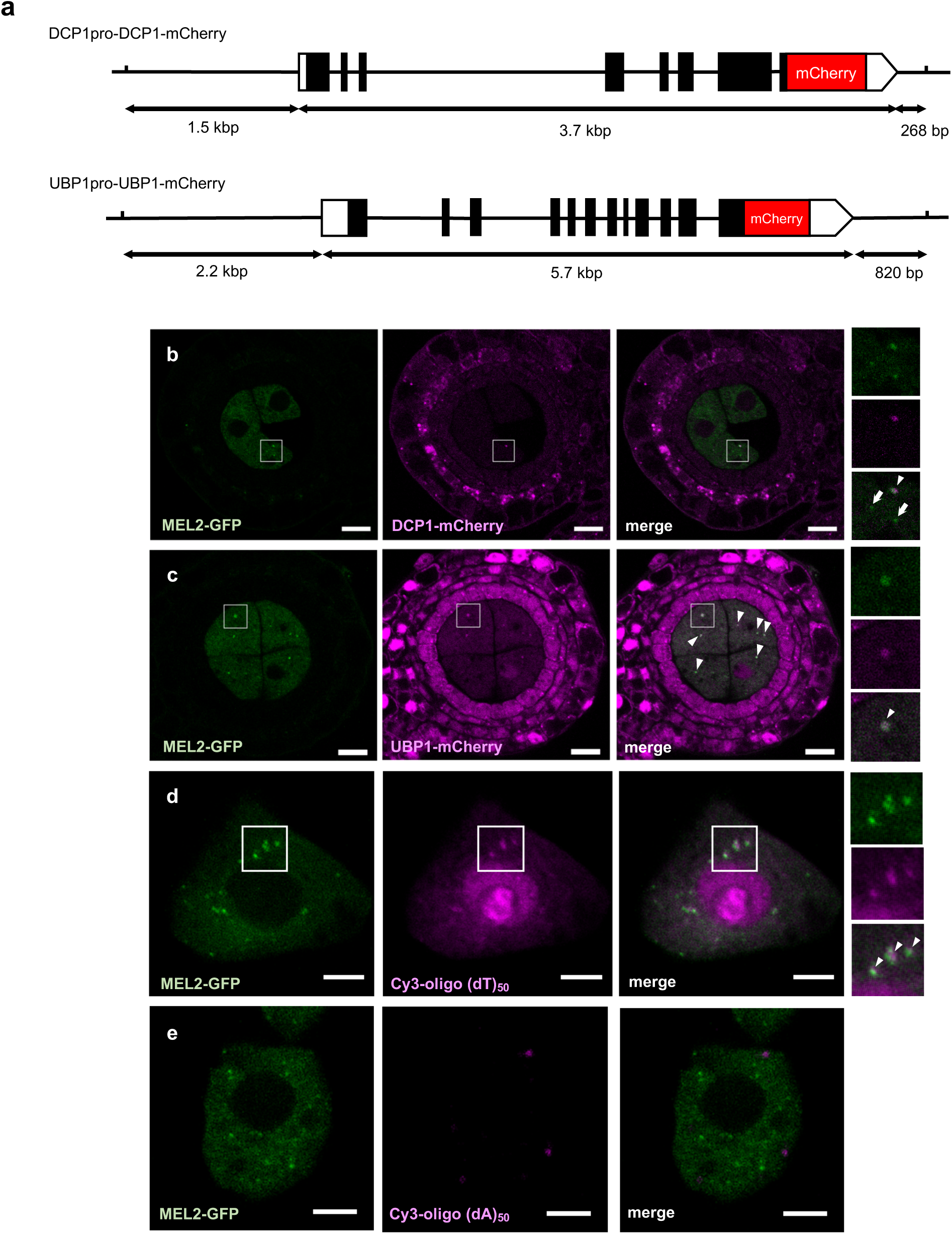
MEL2 granules are associated with P-bodies, stress granules, and poly(A)+ RNA in PMCs. (**a**) Diagram of proDCP1-DCP1-mCherry and proUBP1-UBP1-mCherry transcriptional fusion construct. White, black, and red boxes represent UTRs, exons, and mCherry, respectively. (**b**, **c**) Additional confocal images showing co-localization of MEL2-GFP and DCP1-mCherry (**b**) or UBP1-mCherry (**c**) granules in cross-sections of 0.4-mm (PL stage) anther lobes dissected from transgenic plants expressing both reporter proteins. The rightmost panels show magnified images of the white square regions. Arrowheads indicate co-localized granules. Arrows indicates MEL2-GFP single granules. Bars = 10 μm. (**d**) Additional images of FISH analysis of poly(A)+ RNAs in the MEL2-GFP PMCs showing the association between MEL2 granules and poly(A) RNAs. In some cases, poly(A)+ RNA granules were surrounded by MEL2 granules. PMCs were isolated from 0.4-mm (PL stage) anthers of MEL2-GFP transgenic plants. The rightmost panels show magnified images of the white square regions. White arrowheads indicate co-localized granules. Bars = 5 μm. (**e**) Cy3-oligo(dA)_50_ probe was used as a negative control. Faint foci derived from Cy3 were observed, but these signals were also detected by UV laser and did not overlap with MEL2 granules, indicating they were non-specific signals or distinct from poly(A) RNA granules. Bars = 5 μm.

**Supplementary Figure 6.**
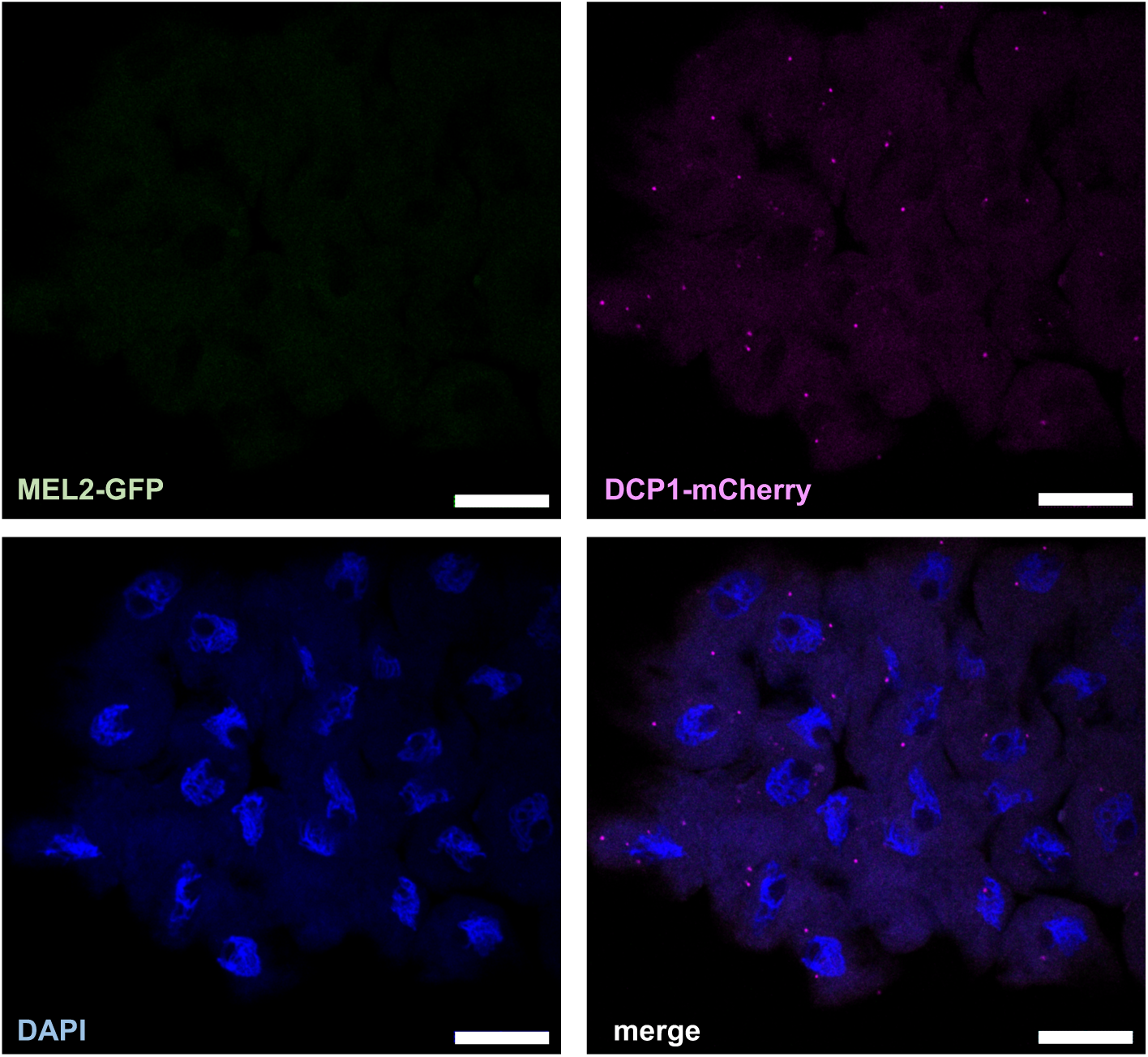
DCP1-mCherry foci in zygotene-pachytene stage PMCs. After MEL2-GFP signals disappeared, P-bodies marked by DCP1-mCherry were still observed after meiotic entry. PMCs were obtained from 0.5-mm ME anthers from MEL2-GFP plants. Bars = 20 μm.

**Supplementary Figure 7.**
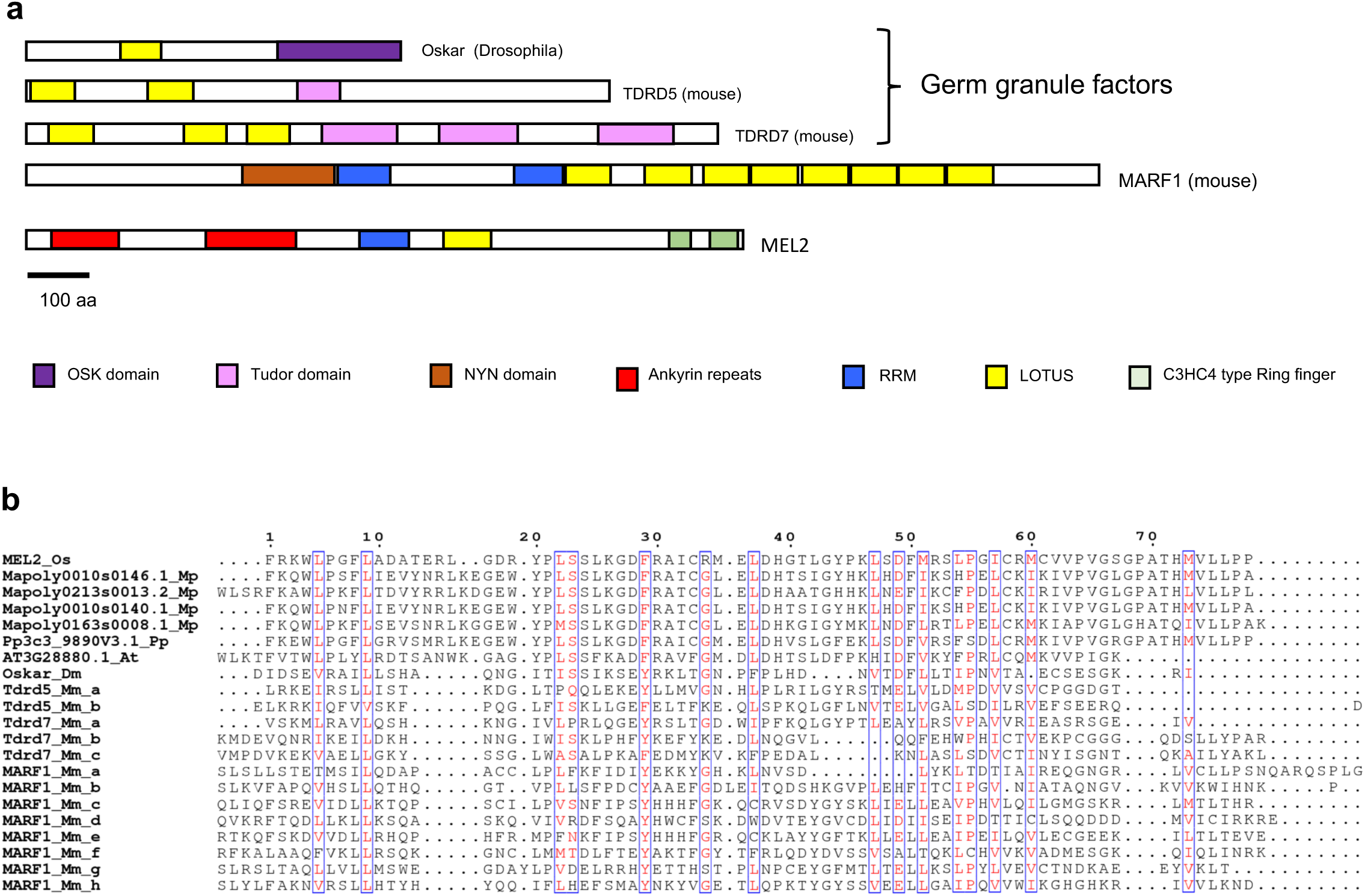
LOTUS domain proteins in animals. (**a**) Schematic representations of domain architecture of LOTUS domain–containing proteins in animals and MEL2 [29]. *Drosophila* Oskar and mouse TDRD5/TDRD7 are germ granule factors in animals. MARF1, which is required for meiotic progression and protection of germline genomic integrity from retrotransposons, contains NYN nuclease domain, two RRMs, and C-terminal repeats of the LOTUS domain. (**b**) Amino acid alignment of LOTUS domains of MEL2-like proteins from plants and LOTUS domain–containing proteins from animals. Os: *Oryza sativa*; Mp: *Marchantia polymorpha*; Pp: *Physcomitrella patens*; At: *Arabidopsis thaliana*; Dm: *Drosophila melanogaster*; Mm: *Mus musculus*.

**Supplementary Figure 8.**
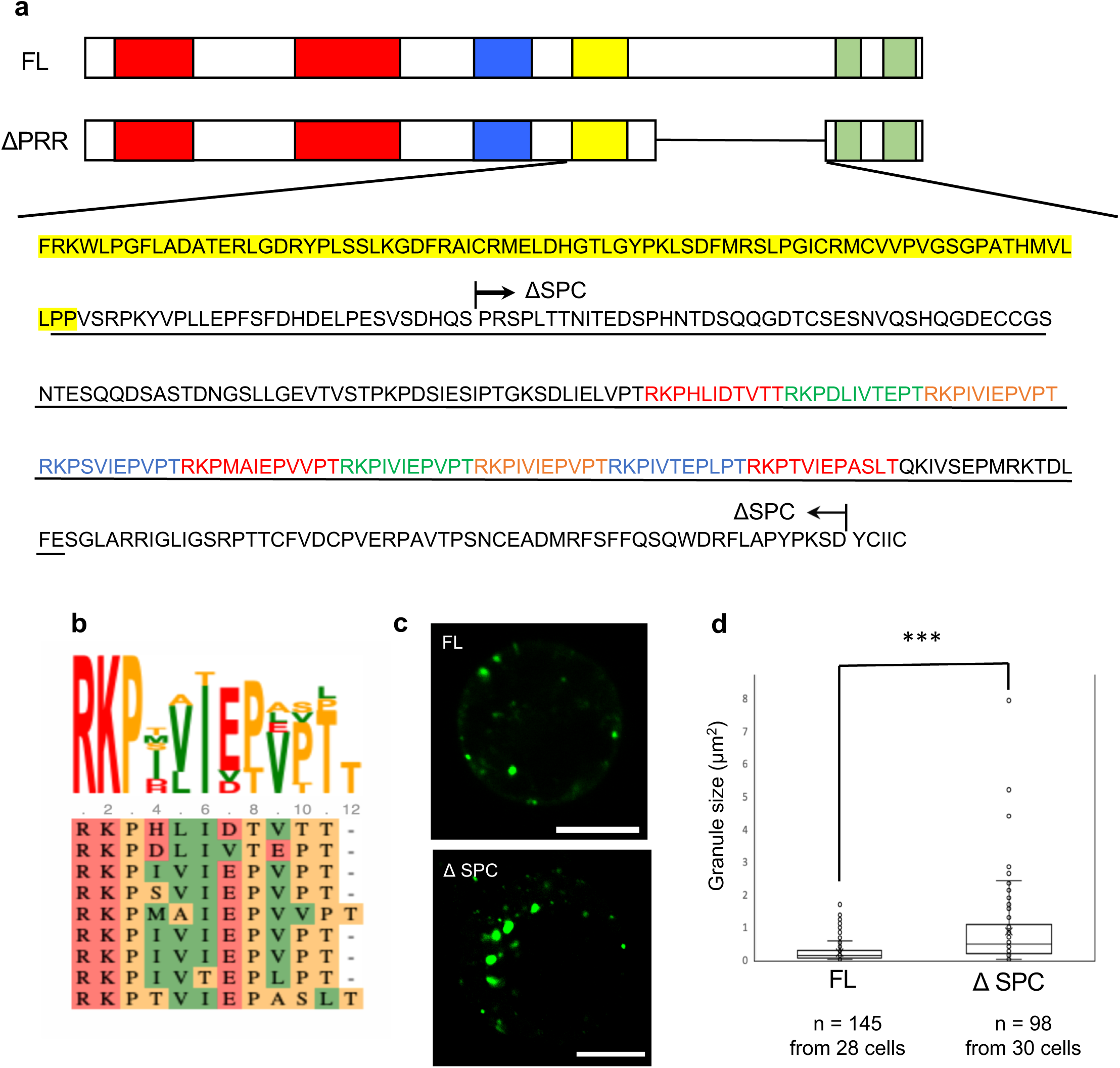
Granule size is regulated by spacer region between LOTUS and RING finger motif. (**a**) Amino acid sequence including LOTUS, IDR-C, and the less-conserved proline rich region between LOTUS and RING finger motif. Highlighted yellow sequence represents the LOTUS domain. Underlined sequence represents the IDR-C region, which contains two proline residues that overlap with the C-terminus of LOTUS. The deleted sequence of ΔPRR is indicated by arrows. Short tandem repeat sequences are shown by colored characters. (**b**) Consensus amino acid of repeat units. (**c**) Representative images of MEL2-GFP and MEL2-ΔSPC-GFP granules in rice protoplast cells. Bars = 10 μm. (**d**) Quantification of granule size in MEL2-GFP– or MEL2-ΔPRR-GFP–expressing cells. Crosses indicate average values of granule size. Triple asterisks indicate a statistically significant difference (*t*-test, P < 0.001).

**Supplementary Figure 9.**
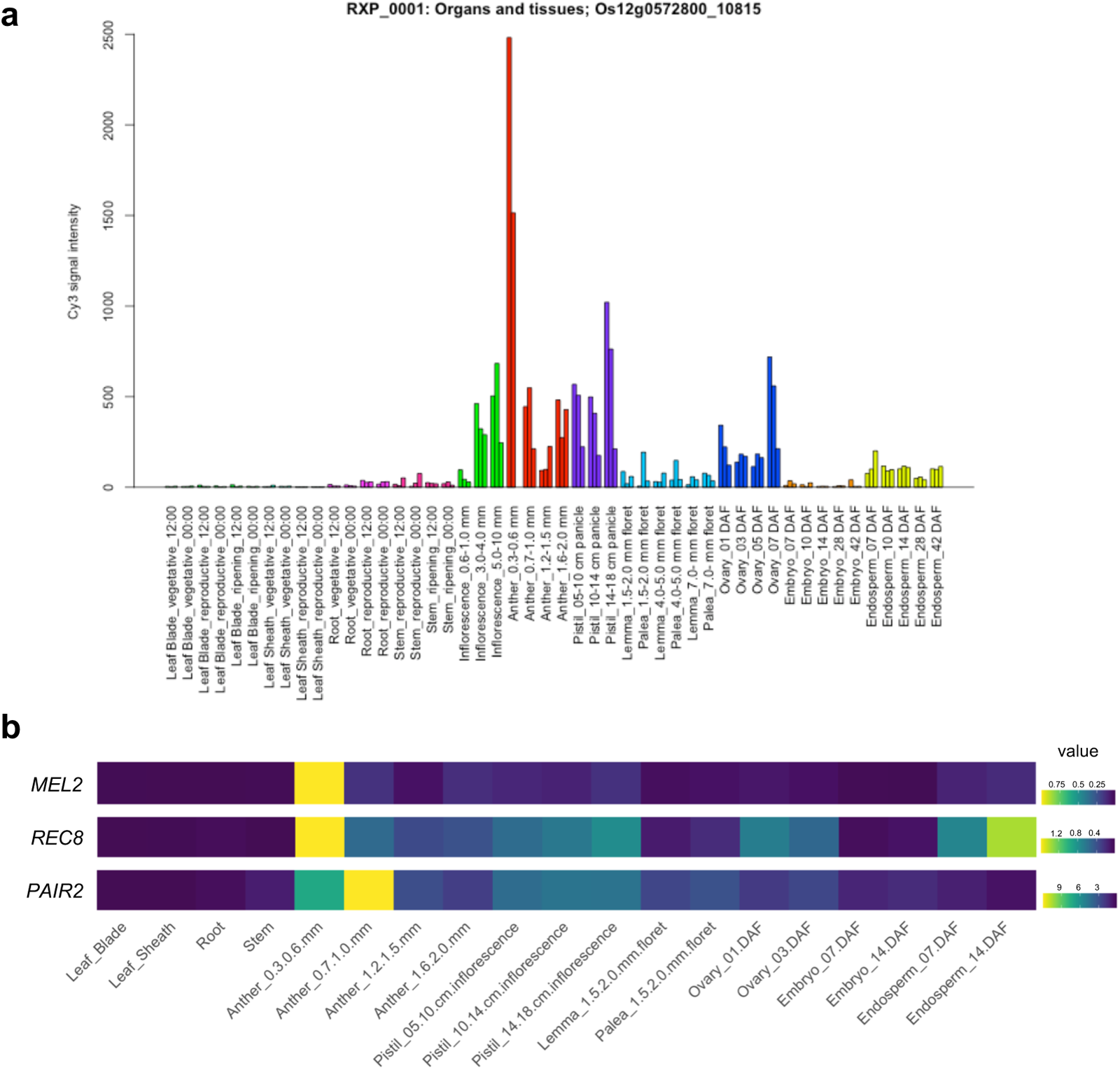
Expression profile of *MEL2* in various tissues and developmental stages. (**a**) Expression profile image obtained from RiceXpro (https://ricexpro.dna.affrc.go.jp). (**b**) Expression heatmap of *MEL2* and meiotic genes *PAIR2* and *REC8*. Normalized and log_2_-transformed values were obtained from a data set deposited at Gene Expression Omnibus (GEO accession: GSE21396) [37]. The values before log_2_ conversion were used to construct the heatmap.

**Supplementary Figure 10.**
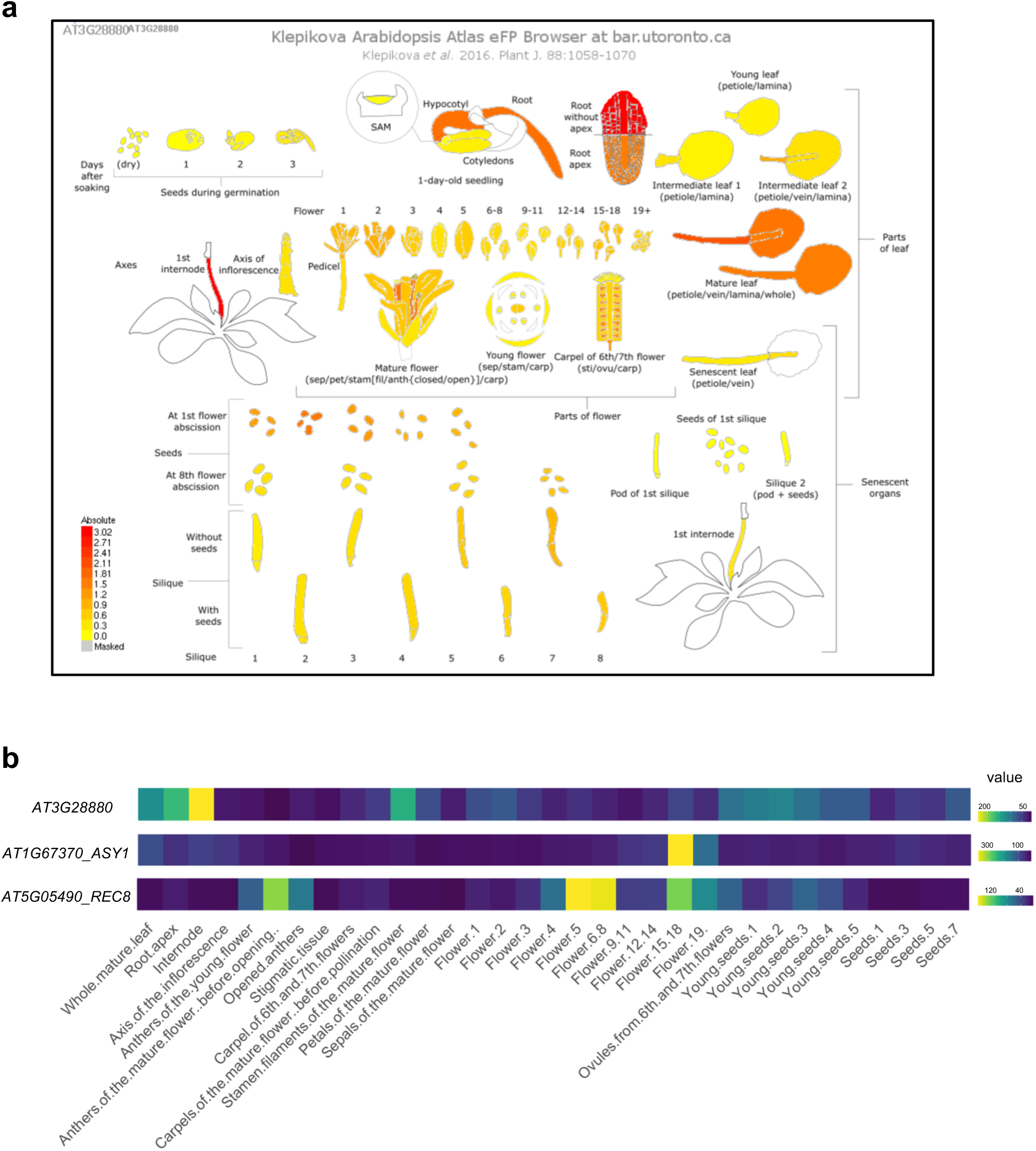
Expression profile of *Arabidopsis* MEL2-like genes in various tissues and developmental stages. (**a**) Image obtained from *Arabidopsis* eFP Browser (http://bar.utoronto.ca/efp/cgi-bin/efpWeb.cgi?dataSource=Klepikova_Atlas). (**b**) Heatmap describing expression values of At3G28880, ASY1, and REC8 during flower development and in reproductive organs. The values were obtained from the TraVA database (http://travadb.org) [38].

**Supplementary Table 1.** Anter developmental stages defined in this study.

| Supplementary Table 1. Anther developmental stages defined in this study. |  |  |  |  |  |
| --- | --- | --- | --- | --- | --- |
| Stages | Floret length (mm) | Anther length (mm) | Germ cell stage | Anther wall stage | Corresponding stage in Ono et al, (2018) [70] |
| Pre-meiosis early (PE) | 0.8-1.2 | 0.2-0.3 | Premeiotic G1-S phase | Three-layered |  |
| Pre-meiosis late (PL) | 1.3-1.7 | 0.3-0.4 | Premeiotic S-G2 phase | Transition from three- to four-layered | ST.1 |
| Meiosis early (ME) | 1.8-2.0 | 0.4-0.5 | leptoten and zygotene | Undifferentiated tapetum and middle layer | ST.2 |

